# Calcium-Regulated Mitochondria Remodeling by Myo19 is Required for Filopodia Tip-Extension

**DOI:** 10.1101/2025.10.04.680418

**Authors:** Sijie Li, I Shenyer Boris, Arnon Henn

## Abstract

Mitochondria are highly adaptable organelles that change their shape, distribution, and movement to meet cellular needs. We investigate how EGF stimulation remodels mitochondrial behavior in human A431 carcinoma cells, where EGF suppresses proliferation but enhances calcium signaling and cell motility. EGF triggers mitochondrial fission, resulting in smaller, more mobile mitochondria that relocate to the tips of newly formed filopodia. Mitochondrial transport involves Kif5B (microtubule-based) and Myo19 (actin-based) motor proteins, which are regulated differently by calcium. EGF stimulation increases cytosolic calcium, weakening Kif5B-mediated transport and making Myo19 the primary transporter. Mitochondrial calcium uptake through the MCU channel is essential for redistribution, shape changes, and filopodia extension. Disrupting mitochondrial fission or calcium buffering impairs mitochondrial motility and the formation of filopodia. High-resolution imaging reveals the coordinated transport of mitochondria, involving fission, motor switching, and localized calcium signaling. We provide insights into how mitochondrial transport supports actin remodeling and protrusive activity, with implications for cell migration and therapies targeting mitochondrial-cytoskeletal interactions.

**HIGHLIGHTS:**

- EGF stimulation remodels the morphology of the mitochondrial network
- Kif5B and Myo19 are required for mitochondrial motility
- Mitochondria display two different velocities depending on the cell region
- Calcium homeostasis contributes to mitochondrial network remodeling

**eTOC BLURB:** Cell powerhouses called mitochondria can change shape and move within cells along tracks called the cytoskeleton. Stimulation of cells by stressors, such as growth factors, can generate signals that cause mitochondria to migrate to the cell edges using molecular motors. Importantly, this movement enables cells to form finger-like projections, providing the energy necessary for cell migration.

## Introduction

Mitochondria are adaptable organelles that regulate their morphology, movement, and positioning in response to cellular needs^1–3^. Besides producing ATP, they regulate calcium homeostasis, generate ROS signaling, facilitate cellular communication, and promote programmed cell death^4–6^. This versatility relies on their ability to dynamically alter their network, connectivity, form, size, and internal structure^7–10^. Signaling pathways regulate mitochondrial dynamics, including the balance of fission and fusion, ER contact sites, cytoskeletal motility, and the activity of its motor proteins^11–14^. Remodeling the mitochondrial network consumes energy and is therefore tightly regulated^15^. Importantly, various stresses and signals, including increased ROS, low oxygen, nutrient shortages, growth factor stimulation (e.g., epidermal growth factor, EGF), and mitochondrial quality control, impact the network’s morphology, motility, and positioning^16–21^. The cytoskeletal filaments, including microtubules and actin, along with their associated motor proteins, play an essential role in coordinating mitochondrial morphology, directing its movement, and positioning^12,22,23^. Highly motile mitochondria are typically found at the leading edge of metastatic breast cancer cells, such as MDA-MB-231 and MDA-MB-436^24,25^. Gaining insight into how cellular physiology drives mitochondrial network remodeling at the molecular level advances our understanding of human health and disease.

Calcium is a key regulator of mitochondrial network remodeling, acting as a signaling molecule in metabolism and apoptosis^26,27^. Calcium enters the mitochondria from specialized ER-mitochondria contact sites marked by Inositol 1,4,5-trisphosphate receptor (IP3R) located on the ER. The release of Ca^2+^ from the ER forms high-concentration "nanodomains" that juxtapose the mitochondrial Ca^2+^ uniporter (MCU), facilitating Ca^2+^ uptake into the mitochondrial matrix^28–32^, VDAC transports cytosolic calcium to the intermembrane space^33^. Calcium exits from the matrix via two IMM channels, the Ca^2+^/Na^+^ exchanger NCLX and the Ca^2+^/H^+^ exchange LETM1^34,35^. The mPTP, a non-selective mitochondrial permeability transition pore channel in the IMM, opens at high calcium levels, releasing proapoptotic proteins and causing depolarization and cell death^36^. NCLX prevents mPTP opening; however, prolonged activation of the mPTP by Prolonged high calcium levels impairs NCLX function^37,38^. Calcium also regulates mitochondrial motor adapter proteins, such as Miro1/2-TRAK1/2 complexes, which interact with Kif5B microtubule-based motor, and Myo19, an actin-based motor, driving mitochondrial motility^39^. Notably, calcium fluctuations in the mitochondrial microenvironment impact its motility, extending beyond its role in mitochondrial fission^40–42^.

Myo19 is an actin-based mammalian myosin motor anchored to the Mitochondrial Membrane (OMM), playing a key role in many aspects of mitochondrial biology. Myo19 is the shortest member of the myosin family, composed of 970 amino acids, with 830 forming the motor and lever arm. Residues 830 to 970 contain OMM and potential Miro and MICOS binding regions^43–45^. Myo19 affects mitochondrial biology by regulating symmetric partitioning during cell division, transporting mitochondria to stress-induced filopodia, tethering mitochondria to ER-associated actin to promote fission, cooperating with Kif5B to clear damaged mitochondria via mitocytosis, and facilitating the localization of MICOS complex-enriched mitochondria in filopodia during single-cell movement^22,43,46–48^. However, the molecular details of Myo19’s role in crista structure and its interaction with Miro proteins are still unclear^45,49^. The mechanism by which Myo19 is activated and regulated remains poorly understood, especially how its motor activity is controlled by its unique enzymatic features that enable it to function both as a transporter and a crosslinker^22,50^.

A suitable model for studying mitochondrial dynamics under calcium regulation is the human epidermoid carcinoma A431 cells. Unlike most cell types, where EGF promotes proliferation, A431 cells exhibit growth-inhibitory responses to EGF at physiological concentrations^51^. EGF causes cell cycle arrest in A431 human epidermoid carcinoma cells and triggers rapid changes in intracellular calcium homeostasis^52–54^. An increase in cytosolic free calcium induced by EGF in A431 cells significantly boosts cell motility and membrane dynamics by enhancing endocytosis^55^. Given our discovery of stress-induced filopodia formation and its dependence on Myo19-mitochondria, this system is highly relevant for studying the molecular coupling between EGF stimulation, calcium homeostasis, and mitochondrial network remodeling.

Here, we elucidate how EGF stimulation triggers a series of events that cause significant changes in mitochondrial shape, movement, and positioning, which are linked to two mitochondrial molecular motors. We have developed a quantitative model based on these observations, which describes how mitochondrial dynamics and movement toward filopodia respond to EGF stimulation in A431 cells. 3D cytoskeletal imaging reveals that our model accurately reflects actin-based filopodia formation, with minimal involvement of microtubules. The movement of mitochondria to the leading edge and filopodia involves both microtubule- and actin-based motors, Kif5B and Myo19, working together. Inside EGF stimulation filopodia, actin filaments and Myo19 are essential for moving mitochondria to the tips of filopodia. Interestingly, calcium buffering affects how Kif5b associates with mitochondria, but it does not significantly influence Myo19’s association with mitochondria *in cells* or *in vitro*. Our findings support the idea that coordinated mitochondrial movement depends on the coordinated action of actin- and microtubule-based motors.

## Results

### Mitochondria translocate to filopodia upon EGF stimulation

High concentrations of EGF stimulation (30 ng/mL) in the A431 epidermal cell line exhibit an epithelial-like morphology in culture, characterized by polarity, resulting in filopodia formation with a high content of Myo19-mitochondria at the leading edge and cellular protrusions (Figure 1A). The EGF stimulation results in a co-movement of ectopically expressed Myo19-GFP and mitochondria, which are highly dynamic, both toward and within filopodia (Movie S1). After EGF stimulation, the length of filopodia increased from 0.9 ± 0.1 µm to 1.9 ± 0.1 µm (n= 100, p < 0.0001), and the average number of filopodia per cell increased from 1.6 ± 0.2 to 11.9 ± 0.5 (n= 100, p < 0.001) (Figure 1B, C). To examine the behavior of the mitochondrial mass at filopodia upon EGF stimulation, we analyzed aging, membrane potential, and mtDNA-dependent mitochondrial activity in EGF-stimulated filopodia, demonstrating mitochondrial viability and the importance of mtDNA presence for filopodia length (Figure S1). Within the filopodia, mitochondria with notably smaller diameters compared to those in the cell body were observed (Figure S2). Under EGF stimulation, the mitochondrial network was qualitatively reshaped into smaller, more mobile structures that translocated to the filopodia. We observed comprehensive mitochondrial remodeling, characterized by significant changes in network dynamics, morphology, and its localization (Figure 1D-G). We categorized mitochondrial locations into four groups. Mitochondria dense network (MDN) is closest to the nucleus, associated with ER, leading edge mitochondria (LE) near the plasma membrane, intermediate mitochondria filopodia within filopodia shaft (IF), and mitochondria at the tip of filopodia (TF, Figure 1D). Quantifying mitochondrial remodeling and morphology in wildtype cells reveals that 87.7% of cells in the wildtype group exhibit MDN localization, while 12.3% show mitochondrial translocation to the LE region. Upon EGF stimulation, 27.5% of cells retained MDN localization, 37.9% had translocation to the plasma membrane (MDN), 7.1% had mitochondrial localization at the IF region, and 27.5% at the TF region (Figure 1E). Our quantitative model strongly suggests that filopodia formation and mitochondrial dynamics are linked to EGF stimulation, resulting in a new distribution of mitochondria within the filopodia shaft and tips.

**Figure 1.**
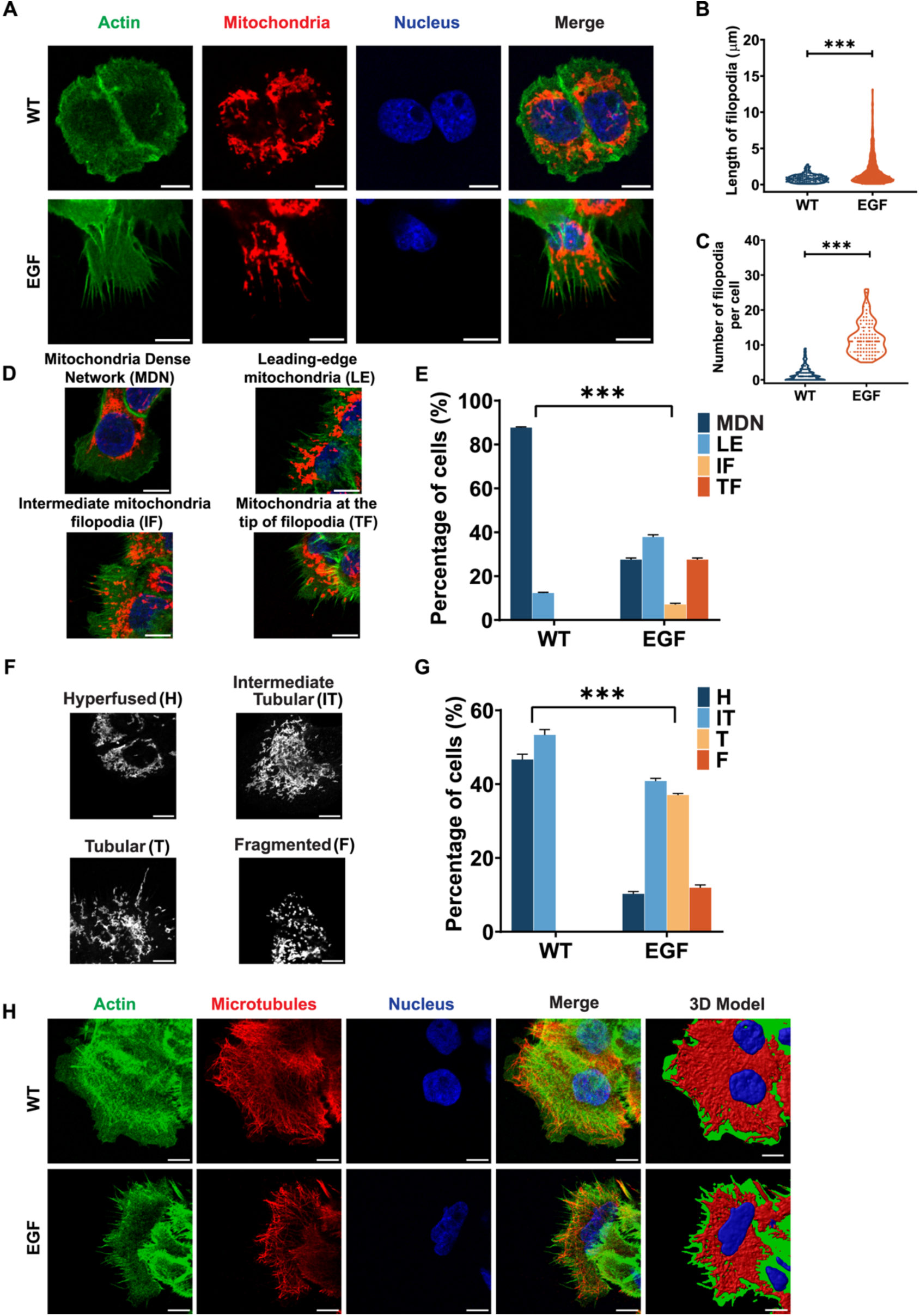
EGF stimulates the formation of filopodia and remodels the mitochondrial network. **(A)** Confocal fluorescence microscopy of A431 cells before (upper panel) and after EGF stimulation (30 ng/ml, lower panel). Cells were serum-starved for 16 h before stimulation and stained for F-actin (green, phalloidin-488), mitochondria (red, α-Tom20 with Alexa Fluor 647-conjugated secondary antibody), and nuclei (blue, Hoechst 33258). Scale bar, 10 μm. **(B and C)** Quantitative analysis of filopodia length **(B)** and number per cell **(C) i**n wildtype (WT) and EGF-stimulated A431 cells (n ≥ 100). Filopodia length measured using the FiloQuant ImageJ plugin (Jacquemet et al., 2017). Data represent mean ± SEM. ***p < 0.001 by unpaired t-test. **(D)** Mitochondrial classification according to cellular localization. Four categories describe mitochondrial positions in EGF-stimulated A431 cells: MDN (mitochondria in the microtubule network region), LE (mitochondria at the leading edge near the plasma membrane), IF (mitochondria in filopodia but not at their tips), and TF (mitochondria at the filopodia tips). Scale bar, 10 μm. **(E)** Quantitative analysis of mitochondrial localization in wildtype (WT) and EGF-stimulated A431 cells (three independent experiments, n ≥ 120). **(F)** Mitochondrial network classification according to morphological dynamics: Hyperfused (H, mitochondrial network in perinuclear region), Intermediate Tubular (IT, mitochondria separated from network into tubular particles), Tubular (T, mitochondria undergoing fission into tubular structures), and Fragmented (F, small mitochondria distributed throughout cells). Scale bar, 10 μm. **(G)** Quantitative analysis of mitochondrial network morphology in wildtype (WT) and EGF-stimulated A431 cells (three independent experiments, n ≥ 120). **(H)** Three-dimensional reconstruction of cytoskeletal structures showing microtubules (α-tubulin with Alexa Fluor 680-conjugated secondary antibody) and F-actin (phalloidin-488) in wildtype (WT, upper panel) and EGF-stimulated (lower panel) A431 cells. Scale bar, 10 μm.

The remodeling of the entire mitochondrial network stems from its dynamic nature; we conducted a thorough quantitative analysis of the morphology and dynamics of the entire mitochondrial mass in response to EGF stimulation. Mitochondrial morphology was classified into four distinct types: hyperfused (H), intermediate tubular (IT), tubular (T), and fragmented (F) (Figure 1F). In the group of wildtype cells, a significant portion (46.7%) showed hyperfused mitochondria, while 53.3% exhibited intermediate tubular mitochondria. Importantly, none of the cells in this group displayed tubular or fragmented mitochondria. However, a notable shift in mitochondrial morphology was observed following EGF stimulation. The percentage of cells with hyperfused mitochondria dropped to 10.3%, while intermediate tubular mitochondria appeared in 40.8% of the cells. Tubular mitochondria were present in 37.0% of the cells, and fragmented mitochondria were observed in 11.9% (Figure 1G). We concluded that EGF stimulation caused significant changes in mitochondrial morphology and dynamics. The transition from mainly hyperfused and intermediate tubular mitochondria in wildtype cells to tubular and fragmented mitochondria after EGF stimulation suggests a complex remodeling process. The appearance of tubular and fragmented mitochondria, which are absent in the wildtype condition, indicates a preference for mitochondrial fission during EGF stimulation.

Cell protrusions form through various mechanisms involving microtubules and actin filaments, with each playing a distinct role in this process. Microtubules can activate Rac1 and generate lamellipodia, providing structural support to filopodia^56^. Imaging of EGF-stimulated cells (Figure 1H) showed that most filopodia lack microtubules. Quantification revealed that in wildtype cells, 53.8% of filopodia had no microtubules, and during EGF stimulation, this number increased to 65.5% (Figures S3A and S3B). Additionally, mitochondria (white) were observed within filopodia (actin, green) without microtubules (Figure S3C). Fluorescence intensity analysis (Figure S3D) confirmed mitochondrial and actin signals in filopodia, but not microtubule signals, indicating that mitochondrial movement in these protrusions depends on actin, not microtubules, during EGF stimulation.

### Mitochondrial fission affects mitochondrial translocation in response to EGF stimulation

To investigate whether mitochondrial fission is essential for filopodia formation and elongation, we treated cells with mdivi-1, followed by EGF stimulation (30 minutes) to measure mitochondrial translocation to filopodia^57^. Mdivi-1 treatment prevented the mitochondria in EGF-stimulated cells from reaching the leading edge and the filopodia, similar to wildtype cells (Figure 2A). Filopodia length in wildtype cells averaged 1.0 ± 0.1 μm, increasing to 2.3 ± 0.1 μm with EGF stimulation (Figure 2B). Mdivi-1 treatment reduced filopodia length to 0.8 ± 0.1 μm, with EGF stimulation showing no significant increase (0.9 ± 0.1 μm). Wildtype cells had 2.4 ± 0.6 filopodia per cell, rising to 11.5 ± 1.5 with EGF stimulation (Figure 2C). Mdivi-1 inhibited filopodia formation, maintaining numbers at 2.0 ± 0.5 and 2.0 ± 0.6 per cell in WT-mdivi-1 and EGF-mdivi-1 groups (Figure 2C). These findings suggest mitochondrial fission is necessary for filopodia formation and elongation. Mdivi-1 possesses a secondary effect of reversible inhibition of mitochondrial Complex 1. However, the Drp1 antagonist behavior serves here as a valuable tool to pinpoint the importance of mitochondrial fission^58^. We then used live-cell imaging to capture fission events further in the leading edge of the cell (Movie S2). We identified two remarkable features that mitochondria exhibit before entering filopodia: aligning vectorially towards, and parallel to, the actin protrusions and a fission event that produces two smaller mitochondria. The latter event enables a single mitochondrion to exhibit high motility towards the tips of filopodia, while mitochondria that did not undergo fission remain stalled. This significant observation demonstrates the production of a smaller mitochondrial mass to enable entry into the filopodia shaft.

**Figure 2.**
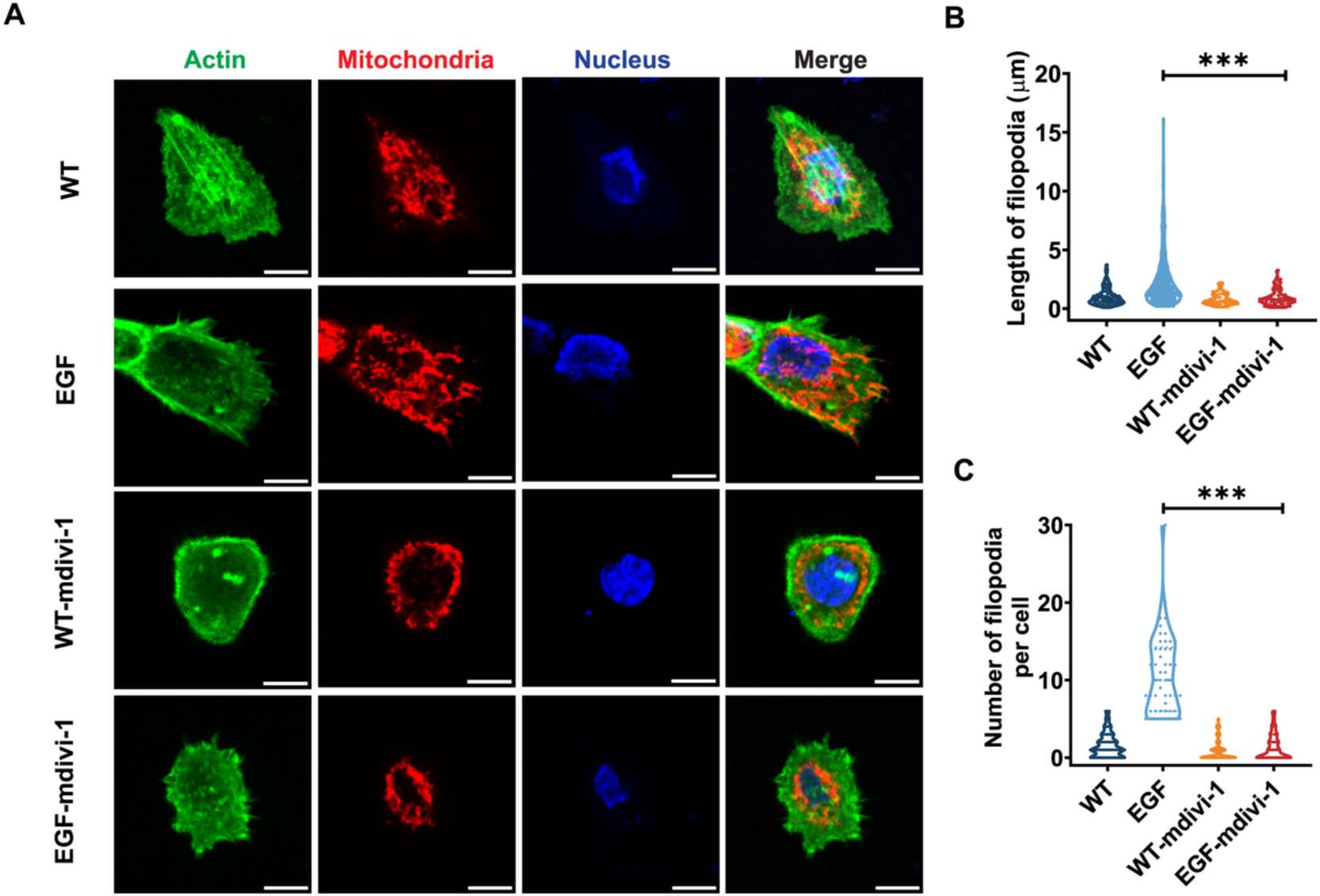
Drp1 activity is required for mitochondrial positioning within filopodia. **(A)** Confocal fluorescence microscopy of A431 cells stained for F-actin (green), mitochondria (red), and nuclei (blue). Four experimental conditions are shown: wildtype (WT), EGF stimulation, WT treated with Mdivi-1 (6 h), and EGF stimulation cells pretreated with Mdivi-1. Mdivi-1 pretreatment (6 h) followed by EGF stimulation (30 min). Merged images demonstrate mitochondrial localization within actin-rich filopodia. Scale bar, 10 μm. **(B)** Quantification of filopodia length across experimental conditions. EGF stimulation significantly increases filopodia length compared to control, while Mdivi-1 treatment blocks EGF-dependent filopodia elongation. Data presented as violin plots (n ≥ 50 cells per treatment). Statistical analysis by one-way ANOVA; ***p < 0.001. **(C)** Quantification of filopodia number per cell. EGF stimulation dramatically increases filopodia formation, which is completely abolished by pretreatment with Mdivi-1. Data presented as violin plots (n ≥ 50 cells per treatment). Statistical analysis by one-way ANOVA; ***p < 0.001.

### Myo19-mitochondrial motility and velocity in the cell body and filopodia indicate active transport

The motility of Myo19-mitochondria toward the cell’s leading edge and filopodia tips is biphasic, featuring two distinct movement patterns followed by a more complex intrafilopodial movement that closely links to the extension and retraction of actin protrusions (Movie S3). The average velocity of mitochondria in the cell body was 211 ± 10 nm/s with a maximum average velocity of 533 ± 27 nm/s (Figure 3B and 3C), which exceeds the velocity of Myo19 (Figure 3D and E) but is lower than the *in vitro* Kif5B velocity^50,59^. Analysis of Myo19-mitochondria movement within filopodia revealed anterograde motion (average velocity = 92 ± 30 nm/s, maximum velocity = 230 ± 70 nm/s, Figure 3D and 3E) and retrograde motion (average velocity = 70 ± 20 nm/s, maximum velocity = 130 ± 30 nm/s, Figure 3F and 3G). We propose that the observed retrograde velocity relates to the natural collapse of filopodia extension caused by actin filament breakage when filopodia reach their extension limit. The retrograde velocity in this motor system is very similar to the actin retrograde velocities reported in filopodia. The maximum anterograde velocity matches the *in vitro* Myo19 actin gliding motility^60^. However, the average velocities are lower, likely due to anchoring interactions, higher loads, and pauses related to filopodia growth dynamics and limited ATP, as previously predicted by our *in vitro* kinetic model and filopodia mechanics to be discussed further^61^. Interestingly, mitochondria in the cell body show more heterogeneous velocities, while actin-rich protrusions display more uniform motility. This motility difference may indicate that the cell body experiences more opposing forces (load), contains longer mitochondria, and has a denser network. We also examined how mitochondrial motility couples with elongation by tracking the filopodia marker FMNL3 during EGF stimulation (Figure S7, Movie S4). The kymographs support that mitochondrial movement in filopodia correlates with filopodial elongation and retraction, suggesting that Myo19-driven mitochondrial motility is linked to actin filament polymerization and depolymerization (Figure S7). Ultimately, under EGF stimulation, Myo19 demonstrates adaptive mitochondrial movement in response to the dynamic behavior of filopodia.

**Figure 3.**
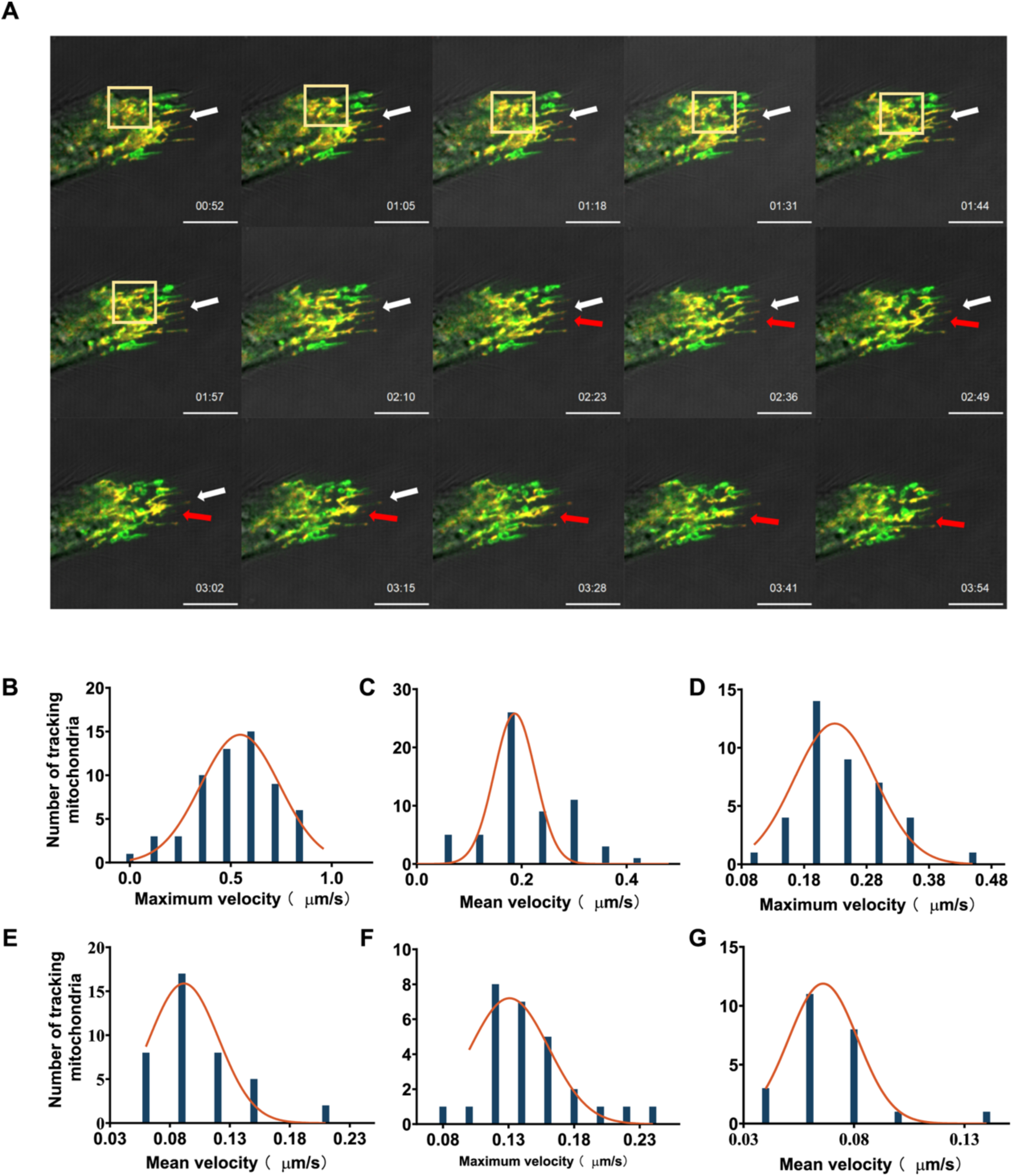
Live-cell imaging reveals Myo19-mediated mitochondrial motility patterns in A431 cells stimulated with EGF. **(A)** Representative time-lapse image series of A431 stable cell line expressing Myo19-GFP with TMRE-labeled mitochondria following EGF stimulation. Images captured at ∼13-second intervals show mitochondrial movements within yellow-boxed regions. White arrows indicate anterograde motility (post-fission event); red arrows indicate retrograde motility. Scale bar, 10 μm. **(B and C)** Distribution histograms of maximum **(B)** and mean **(C)** velocities of mitochondrial motility in the cytoplasm upon EGF stimulation (n ≥ 60 mitochondrial movement events tracked). Orange curves represent Gaussian fits to data distributions. **(D and E)** Maximum **(D)** and mean **(E)** velocity distributions of anterograde Myo19-mitochondria transport within filopodia (n = 40 mitochondrial movement events tracked). **(F and G)** Maximum **(F)** and mean **(G)** velocity distributions of retrograde Myo19-mitochondria transport within filopodia (n = 28 mitochondrial movement events tracked). All data were acquired by time-lapse fluorescence microscopy at 1-second intervals. Particle tracking and velocity analysis were performed using Imaris 8.1 software.

### Kif5B and Myo19 cooperate to enable mitochondria to translocate to filopodia

Myo19 mediates mitochondrial motility in actin-based filopodia structures, while Kif5B is the primary molecular motor responsible for mitochondrial motility along microtubules^22,43,62^. To clarify the coordination and functions of Myo19 and Kif5B in mitochondrial motility during EGF stimulation, we established two cell lines expressing inducible Kif5B and Myo19 shRNAs (Figures S2A and S2B). Kif5B shRNA and Myo19 shRNA reduced protein levels by 50% and 73%, respectively (Figures S2C and S2D). The knockdown cell lines showed slight changes in OMM protein composition (Figures S2D to S2F). We then stimulated filopodia with EGF and observed that knockdowns of Kif5B and Myo19 impaired mitochondrial translocation and decreased filopodia formation, even in the presence of EGF stimulation (Figure 4A). Myo19 knockdown caused a significant reduction in mitochondrial dynamics, favoring fusion, with mitochondria mainly remaining stationary in the MDN. Both knockdown cell lines also exhibited shorter filopodia, indicating that depletion of Kif5B or Myo19 hindered filopodia elongation.

**Figure 4.**
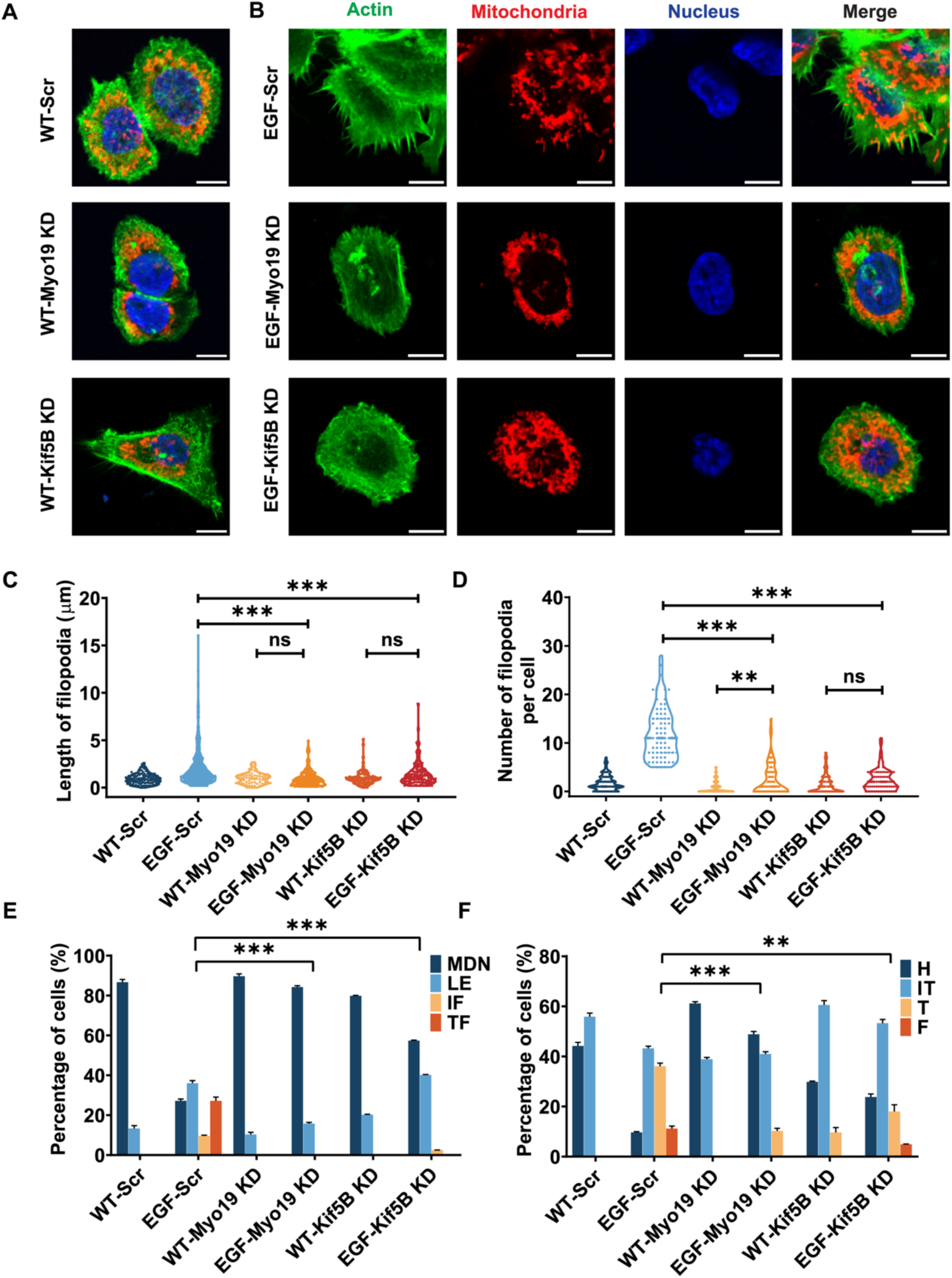
Mitochondrial dynamics and morphology are coupled to filopodia formation in EGF stimulation A431 cells following Myo19 or Kif5B knockdown. (**A and B**) Confocal fluorescence microscopy of A431 cells stained for actin (green), mitochondria (red), and nuclei (blue) in wildtype scrambled control (WT-Scr), Myo19 knockdown (WT-Myo19 KD), and Kif5B knockdown (WT-Kif5B KD) conditions. (A) Cells without EGF stimulation. (B) EGF stimulation cells. Scale bar, 10 μm. **(C and D)** Quantification of filopodia characteristics in EGF stimulation conditions. (C) Filopodia length measurements show that Myo19 and Kif5B knockdown prevent EGF-induced filopodia elongation. (D) Filopodia number per cell analysis demonstrates that EGF stimulation dramatically increases filopodia formation in control cells, which is significantly reduced by Myo19 knockdown and completely abolished by Kif5B knockdown. Data from three independent experiments (n ≥ 120 cells per condition). **(E)** Quantitative analysis of mitochondrial subcellular localization categorized as MDN (mitochondria distributed near nucleus), LE (localized to cell edge), IF (intermediate filaments), and F (filopodia). EGF stimulation promotes mitochondrial redistribution to filopodia tips in control cells, while Myo19 and Kif5B knockdown alter this localization pattern. **(F)** Analysis of mitochondrial morphology classified as H (hyperfused), IT (intermediate tubular), T (tubular), and F (fragmented). EGF stimulation affects mitochondrial morphology, whereas the knockdown of Myo19 and Kif5B reveals distinct morphological profiles. Data represent mean ± SEM from three independent experiments (n ≥ 120 cells per condition). Statistical analysis by one-way ANOVA; ns, not significant; *p < 0.05; **p < 0.01; ***p < 0.001.

We carried out a detailed quantitative analysis of four parameters: filopodia length, the number of filopodia per cell, mitochondrial positioning, and mitochondrial morphology, to assess the effects of Myo19 and Kif5B knockdown with or without EGF stimulation (Table 1). The behaviors observed for Myo19 and Kif5B knockdowns strongly suggest that both are vital for mitochondrial movement in response to EGF.

**Table 1:**
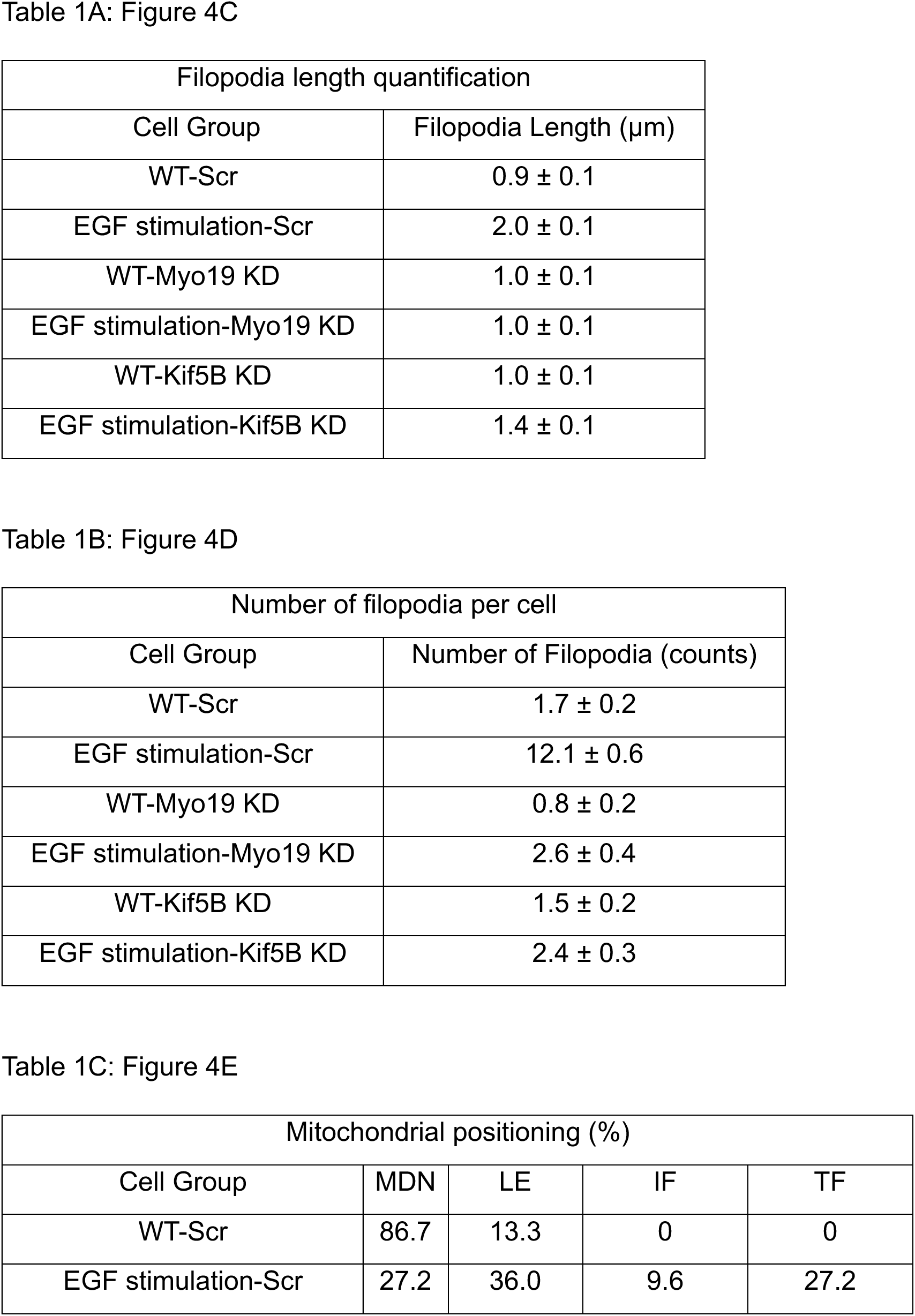
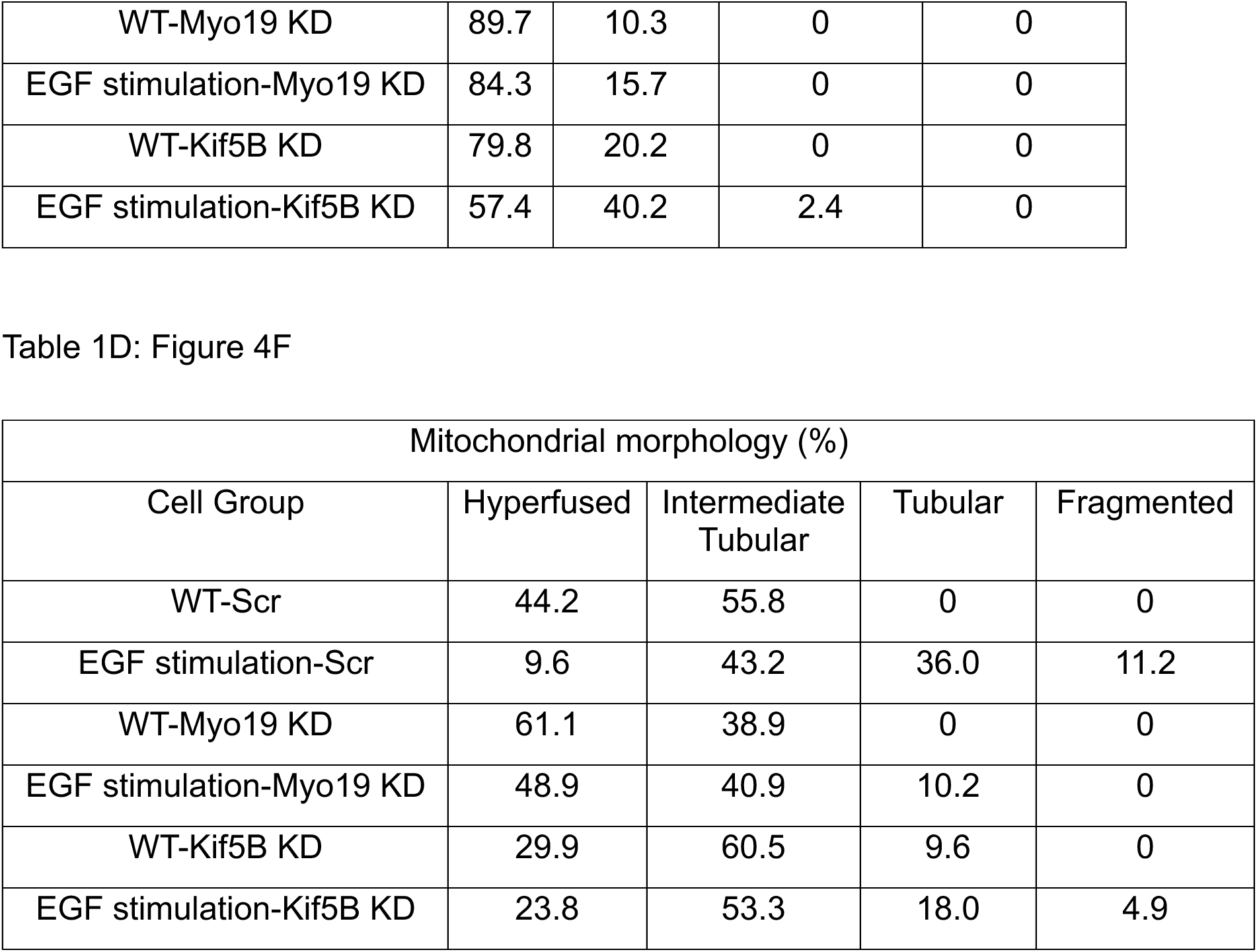
Kif5B and Myo19 cooperate, mitochondria translocate to filopodia.

Specifically, the length and number of filopodia per cell (Table 1A) show a similar decrease for each knockdown. However, the effect of Myo19 knockdown appears more pronounced than that of Kif5B knockdown when compared to EGF-stimulated wildtype cells expressing scrambled shRNA in terms of filopodia length, supporting the idea that Myo19-mitochondria contribute to actin protrusions through a specific function. The number of filopodia did not differ between the two knockdowns (Table 1B), indicating that the actin remodeling signaling responsible for forming actin protrusions, contributed by Myo19 and Kif5B, likely performs similar roles. Next, we quantified mitochondrial positioning and morphology after EGF stimulation in Myo19- and Kif5B-knockdown cell lines (Tables 1C and 1D). The contribution of Myo19 and Kif5B to mitochondrial movement toward the leading edge of cells (LE) is significant, as Kif5B and Myo19 produce comparable levels of motility relative to EGF-stimulated cells with scrambled knockdown, despite Kif5B depletion, and may even be considered compensatory.

Although both are necessary, their impact is not equal. When examining mitochondrial morphology resulting from EGF stimulation, which is regulated by fission-fusion dynamics, Myo19 and Kif5B knockdown cell lines show a significant effect compared to WT. Both are essential for shaping mitochondrial morphology and dynamics through different mechanisms. However, Kif5B knockdown results in more tubular mitochondria than Myo19 knockdown, indicating that Myo19 promotes mitochondrial dynamics toward shorter, more narrow structures that may better facilitate motility into elongated actin protrusions. To verify that Myo19 motor activity is vital for our findings, we performed a rescue experiment using the Myo19 tail domain (Figure S6A-D). The C-terminal domain, which interacts with OMM proteins and contains the Myo19 OMM membrane anchoring domain, was unable to rescue the significant effects seen with full-length Myo19 knockdown, as quantified (Figures S6E-F). Recently, it was shown that Myo19 knockdown does not induce crista deformation in U2OS cell lines^63^.

### Mitochondria’s calcium buffering affects mitochondrial translocation

In A431 cells, high concentrations of EGF (from 26.0 to 54.4 ng/ml) cause an increase in cytoplasmic calcium, with the calcium response lasting from 8 to 32 seconds and reaching its peak between 20 and 410 seconds^64^. The ER is a significant source of calcium release into the cytosol in response to EGF stimulation. However, it remains unclear when the calcium flux begins, when the calcium concentration reaches its peak, and how mitochondria respond to changes in calcium levels. To examine calcium variations in mitochondria upon EGF stimulation, we labeled mitochondrial calcium levels with Mito-R-GeCo1 (Figure 5A, Movie S5 and S6). Mitochondrial calcium levels during EGF stimulation were monitored by overexpressing Mito-R-Geco for live fluorescence microscopy. We tracked calcium changes in A431 cells transiently expressing Mito-R-Geco, added EGF, and then started recording calcium fluctuations every 10 seconds. After 190 seconds, mitochondrial calcium peaked and then declined (Figure 5A).

**Figure 5.**
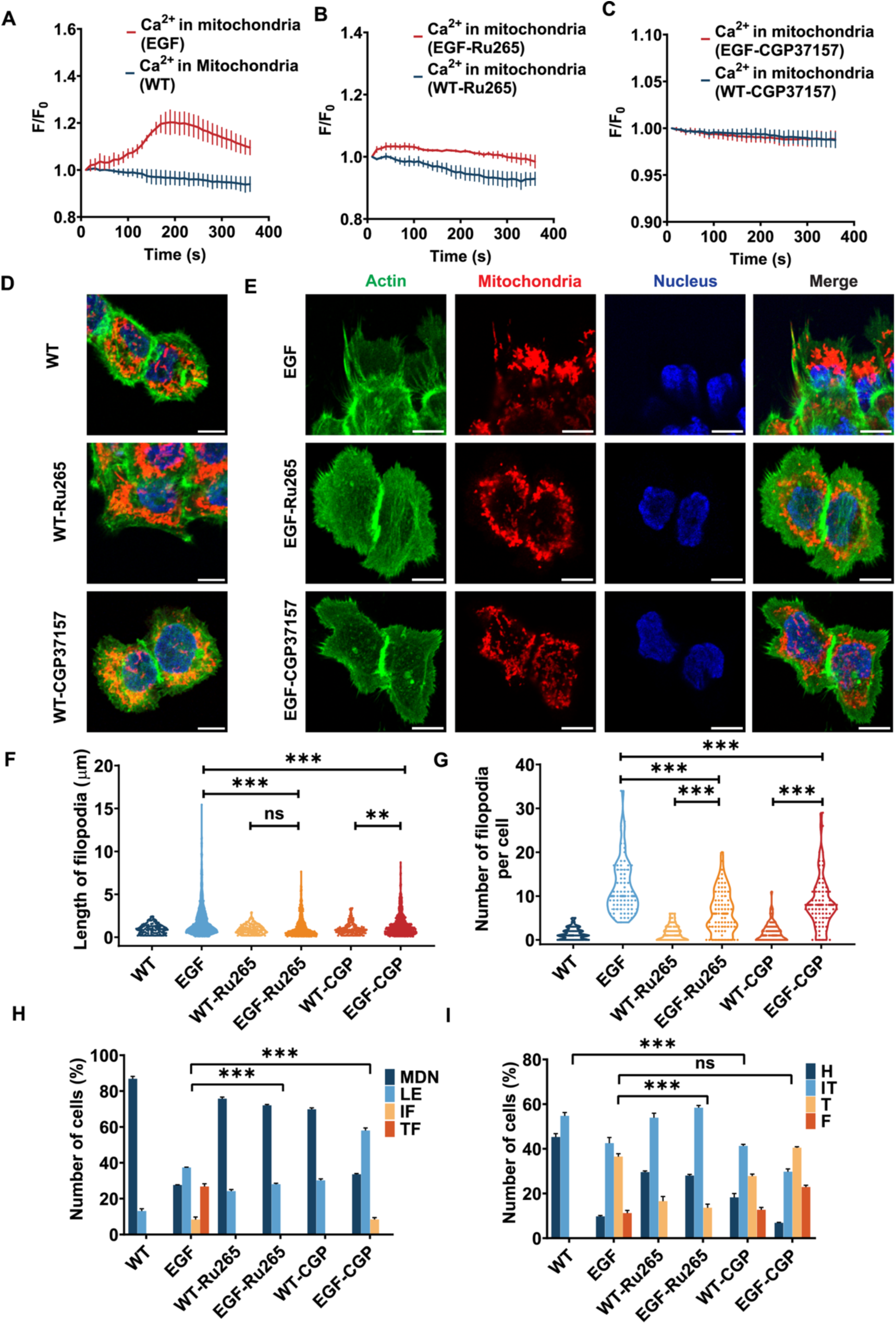
Mitochondrial calcium buffering regulates the formation of filopodia and the localization of mitochondria. **(A-C)** Time-course analysis of mitochondrial Ca²⁺ levels in A431 cells measured by fluorescence intensity (F/F₀) normalized to initial values at 10 s post EGF stimulation. **(A)** EGF stimulation (red) increases mitochondrial Ca²⁺ compared to the untreated control (black). **(B)** Pretreatment with Ru265 (1 μM, mitochondrial calcium uniporter inhibitor) blocks EGF stimulation mitochondrial Ca²⁺ elevation. **(C)** Pretreatment with CGP37157 (10 μM, mitochondrial Na⁺/Ca²⁺ exchanger inhibitor) prevents EGF-induced mitochondrial Ca²⁺ changes. Data represent mean ± SEM. **(D and E)** Confocal fluorescence microscopy of A431 cells stained for actin (green), mitochondria (red), and nuclei (blue). Scale bar, 10 μm. **(D)** Untreated control cells and cells pretreated with Ru265 or CGP37157 without EGF stimulation. (E) Cells were pretreated with inhibitors (10 min) followed by EGF stimulation. **(F and G)** Quantification of filopodia length (F) and number per cell (G). EGF stimulation significantly increases both parameters in control cells, while calcium buffering inhibitors (Ru265 and CGP37157) block these effects. Data from three independent experiments (n ≥ 120 cells per condition). **(H)** Analysis of mitochondrial subcellular localization categorized as MDN (mitochondria distributed in nucleus), LE (localized to cell edge), IF (intermediate filaments), and TF (throughout filopodia). EGF promotes mitochondrial redistribution to filopodia, a process inhibited by calcium-buffering agents. **(I)** Mitochondrial morphology analysis showing distribution according to H (hyperfused), IT (intermediate tubular), T (tubular), and F (fragmented). EGF stimulation alters mitochondrial morphology, with calcium buffering inhibitors affecting this remodeling. Data represent mean ± SEM from three independent experiments (n ≥ 120 cells per condition). Statistical analysis by one-way ANOVA; ns, not significant; **p < 0.01; ***p < 0.001.

Next, we used inhibitors to block calcium flow in the mitochondria to investigate the roles of MCU and NCLX as primary channels for mitochondrial calcium buffering during EGF stimulation. Ru265, a new inhibitor of the mitochondrial MCU channel responsible for calcium uptake into mitochondria, has a structure similar to Ru360, featuring two bridged Ru centers and ammine ligands. Additionally, Ru265 exhibits relatively low toxicity and does not interfere with other mitochondrial functions or calcium fluxes, such as those mediated by the HCX and NCLX channels^65^. For live imaging experiments, A431 cells expressing Mito-R-Geco were treated with Ru265 (1 μM) for 10 minutes, followed by stimulation with EGF. Calcium level changes were monitored using Mito-R-Geco (red) at 10-second intervals. The F/F_0_ value peaked at 1.2 during mitochondrial calcium increase after EGF stimulation (Figure 5B). When Ru265 blocked the MCU, the maximum F/F_0_ ratio was 1.02, indicating that mitochondrial calcium uptake was hindered (Figure 5A), which suggests that the MCU is the primary channel for calcium entry following EGF stimulation. CGP37157 is an inhibitor of the mitochondrial NCLX channel, which regulates calcium release from mitochondria^66^. A431 cells expressing Mito-R-Geco were treated with CGP37157 (10 μM, 10 minutes), followed by EGF stimulation (Figure 5C). Calcium levels were tracked with Mito-R-Geco (red) at 10-second intervals. These results show that when CGP37157 inhibits NCLX, it disrupts calcium exit from the mitochondria. The measurements shown in Figures 5A-C were supported by IF microscopy (Figures 5D-E), which revealed an increase in calcium levels in the mitochondria. Next, we stained the mitochondria with Tom20 and actin with phalloidin, demonstrating that the MCU and NCLX inhibitors Ru265 and CGP37157, respectively, reduced filopodia formation in response to EGF stimulation. CGP37157 also promoted mitochondrial fission, leading to more fragmented mitochondria (Figures 4D and 4E).

Filopodia length is more affected by Ru265 than CPG37157 (Tables 2A & 2B); however, the number of filopodia shows the opposite trend, with the latter nearly reducing the number to that of WT without EGF. These results suggest that calcium uptake regulates filopodia length by influencing the function and targeting of Kif5B, as well as Miro1/2. Calcium spikes, triggered by EGF stimulation, may modulate signaling pathways and activate actin remodeling at the cell’s leading edge^67^. Mitochondrial positioning and shape are strongly impacted differently by calcium uptake inhibition compared to blocking calcium efflux. Inhibiting calcium efflux has a minor effect on the ability to translocate mitochondria to the filopodia shaft, resulting in a slight reduction in reaching the filopodia tips. However, calcium uptake significantly restores mitochondrial positioning to WT levels (Table 2C).

**Table 2:**
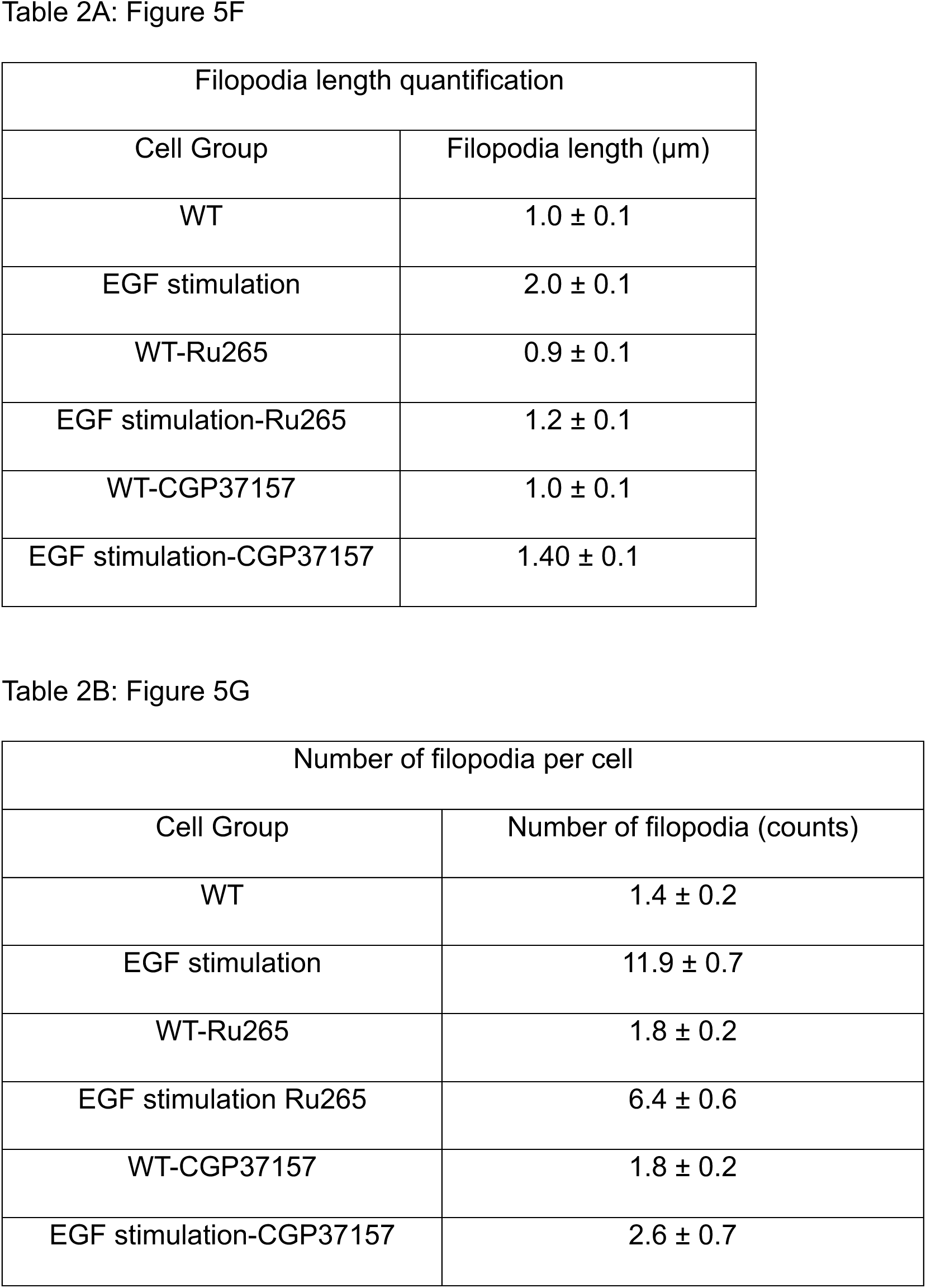
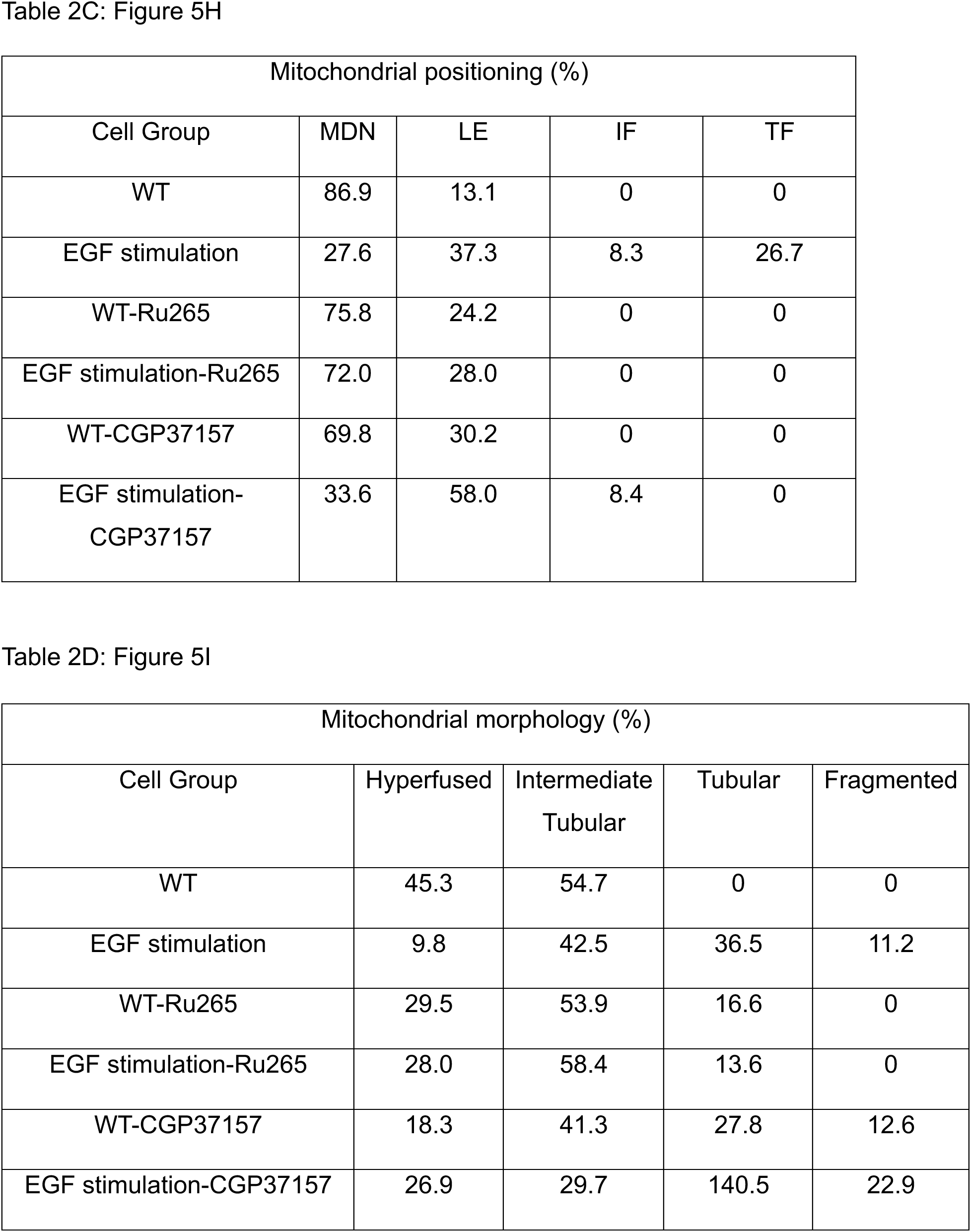
Mitochondria calcium buffering affects mitochondrial translocation.

Furthermore, this finding supports the idea that mitochondrial shape changes during calcium influx rather than calcium efflux. Our results (Table 2D) show that calcium uptake significantly decreases the mitochondria’s tendency to form a smaller network. In contrast, blocking calcium efflux has an even greater effect on network resizing than the WT-EGF stimulation cell group (Table 2D). In summary, CGP37157 treatment increases mitochondrial fission following EGF stimulation, suggesting that this process is triggered by mitochondrial calcium overload. However, calcium uptake was not detected in the calcium curve from direct fluorescence measurements, most likely because the first measurement was taken at 10 seconds, as calcium uptake occurs rapidly. Pretreating cells with CGP37157 under normal conditions before EGF stimulation may already saturate mitochondrial calcium.

### Mitochondrial calcium buffering and release are suggestive of the distribution of mitochondrial motor proteins in cells

Next, we used SR with IF to examine whether the rapid rise in cytosolic calcium influences the colocalization of the molecular motors Kif5B and Myo19 with mitochondria. Figures 6C and 6D display the colocalization of Kif5B with mitochondria, both with and without EGF stimulation, under inhibited MCU conditions. We quantified colocalization in each dataset using the Pearson correlation coefficient (PCC). Notably, we observed a significant decrease in Kif5B-mitochondria colocalization after EGF stimulation (Figure 6E). The PCC in wildtype cells was 0.48 ± 0.01, but it dropped to 0.41 ± 0.01 in EGF stimulation cells (t-test, P < 0.01). However, treating the cells with an MCU inhibitor prevented this decline, resulting in nearly identical Kif5B-mitochondria colocalization with or without EGF stimulation (Figure 6E, WT cells-CGP37157 and WT cells-CGP37157-EGF stimulation; PCCs were 0.50 ± 0.01 and 0.49 ± 0.01, respectively, P > 0.05). These results support the idea that rapid increases in cytosolic calcium promote the delocalization of Kif5B from mitochondria, likely through conformational changes in Miro1/2 induced by calcium binding.

**Figure 6.**
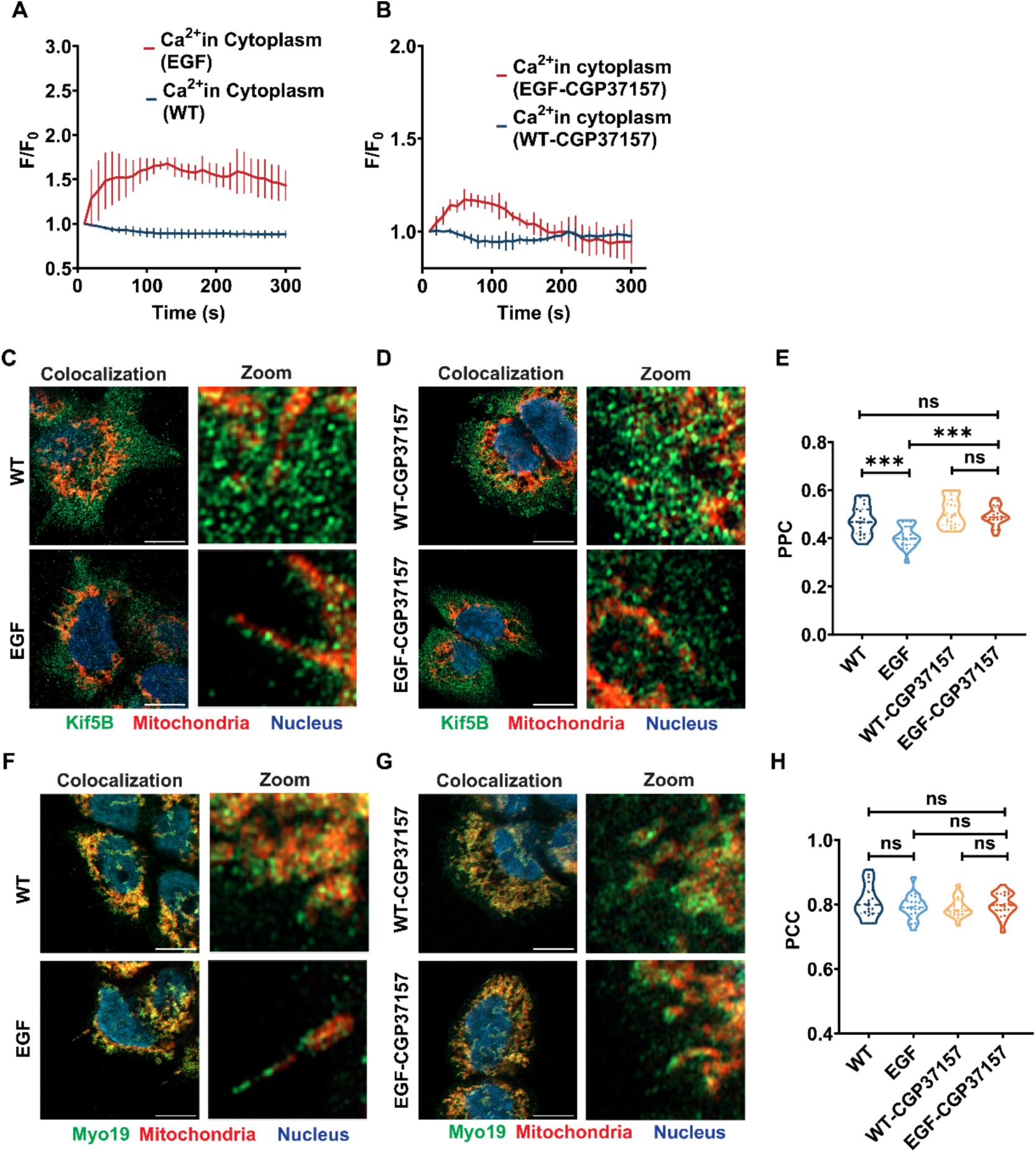
Mitochondrial calcium buffering regulates Kif5B-mitochondria association upon EGF stimulation. **(A and B)** Time-course analysis of cytoplasmic Ca²⁺ levels in A431 cells measured by fluorescence intensity (F/F₀) normalized to initial values at 10 s post EGF stimulation. (A) EGF stimulation (red) significantly elevates cytoplasmic Ca²⁺ levels compared to the untreated control (black). (B) Pretreatment with CGP37157 (10 μM, blue) blocks EGF stimulation-induced cytoplasmic Ca²⁺ elevation (red). Data represent mean ± SEM. **(C and D)** Confocal fluorescence microscopy of A431 cells showing Kif5B (green), mitochondria (red), and nuclei (blue). Scale bar, 10 μm. (C) Untreated control and EGF stimulation cells demonstrating enhanced Kif5B-mitochondria colocalization upon EGF stimulation. (D) Cells pretreated with CGP37157 (10 μM, 10 min) followed by EGF stimulation show reduced colocalization. The right panels show magnified regions. **(E)** Quantification of Kif5B-mitochondria colocalization by Pearson correlation coefficient (PCC). EGF stimulation significantly increases colocalization, which is blocked by pretreatment with CGP37157 (n = 20 cells per condition). **(F and G)** Confocal fluorescence microscopy showing Myo19 (green), mitochondria (red), and nuclei (blue). (F) Untreated control and EGF-stimulated cells. (G) Cells were pretreated with CGP37157 and then stimulated with EGF. The right panels show magnified regions. Scale bar, 10 μm. **(H)** Quantification of Myo19-mitochondria colocalization by PCC shows no significant changes across conditions (n = 20 cells per condition). Data represent mean ± SEM. Statistical analysis by one-way ANOVA; ns, not significant; ***p < 0.001. Images acquired using Zeiss LSM980 confocal microscope. At least two plausible mechanisms by which calcium could regulate Myo19-mitochondria activities involve interactions of the Myo19 tail domain (aa 900-970) with Miro1/2 and a calcium-sensitive light chain that may regulate its motor activity. We conduct parallel experiments on Kif5B-mitochondria to examine the colocalization of Myo19 with mitochondria (Figures 6F and 6G). Importantly, we observed no significant reduction in Kif5B-mitochondria colocalization after EGF stimulation (Figure 6H; PCC in WT cells was 0.81 ± 0.01, decreasing to 0.80 ± 0.01 in EGF stimulation cells, t-test P > 0.05). Treating cells with an MCU inhibitor did not affect the interaction and resulted in nearly identical Myo19-mitochondria colocalization after EGF stimulation (Figure 6H). The values for WT cells-CGP37157 and WT cells-CGP37157-EGF stimulation were similar to those for PPC, with values of 0.8 ± 0.01 and 0.8 ± 0.001, respectively (P > 0.05). These findings indicate that rapid increases in cytosolic calcium promote the colocalization of Kif5B with mitochondria.

## Discussion

EGF stimulation at low concentrations promotes filopodia formation while inhibiting cell proliferation. We have demonstrated a direct link between strong EGF signaling and a rapid calcium buffering response. This response facilitates remodeling of the mitochondrial network, actively moving its mass from the microtubule network in the cell center to the tips of filopodia through the coordinated activities of molecular motors. This process aligns with the remodeling of the mitochondrial network, shifting from a hyperfused to a fragmented state, which enhances the navigation of cellular protrusions. Two key motor proteins coordinate mitochondrial movement: Kif5B (microtubule-based) and Myo19 (actin-based) in A431 epidermal cell lines. Our findings suggest a handover mechanism in which Kif5B initially transports mitochondria along microtubules toward the cell periphery, and then Myo19 takes over for the final movement along actin filaments within filopodia. Part of this process involves the different calcium sensitivities of these motors. Knockdown studies confirm that both proteins are essential for the formation of effective filopodia and the proper positioning of mitochondria.

Mitochondrial calcium buffering plays a vital role in regulating these motor proteins. EGF stimulation triggers calcium signaling pathways, and increased cytosolic calcium causes Kif5B to detach from mitochondria, while Myo19-mitochondria interactions stay intact. This calcium-dependent process facilitates the transfer of transport from microtubules to actin filaments.

### EGF stimulation triggers cellular biochemical pathways to increase mitochondrial motility

EGF stimulation activates EGFR signaling and promotes EGFR translocation to the mitochondria, ER, and nucleus^68,69^. EGFR-mediated signaling causes localized calcium oscillations at contact sites between the plasma membrane and ER, as well as between the ER and mitochondria. This phenomenon is sometimes referred to as noncanonical stress-induced EGFR trafficking and signaling, and it has significant implications for disease progression and therapeutic outcomes. Mitochondria sense calcium signals, which increase metabolism and ATP production, both of which are critical for cortical actin remodeling^71^. In A431 cells, the localization of EGFR to mitochondria regulates autophagy and cell death pathways, linking cell viability to mitochondrial EGFR levels^72^. Our *in-cell* studies show that within 30-45 minutes, mitochondria remain functional, undergo fission, move to the cell periphery, and stay active as part of the long-term EGF response. Initially, mitochondria boost active filopodia dynamics, which are necessary for cell migration. EGF stimulation causes immediate increases in intracellular calcium in human glioma and A431 cells^73^. Mitochondrial EGFR (mitoEGFR) enhances ATP production by promoting oxidative phosphorylation and reducing glucose dependence^74^. Internalized EGFR also decreases mitochondrial fusion by interacting with Mfn1, supporting our observation that mitochondria can move as smaller units^75^. In A431 cells, EGF activates calcium signaling through Phospholipase C (PLC), leading to Store-Operated Calcium Entry (SOCE) and activation of IP3 receptors on ER membranes^76–78^. This series of events causes calcium release from the ER, increasing cytosolic calcium levels and promoting mitochondrial calcium uptake. Our results suggest that the ER plays an important role in maintaining cytosolic calcium levels, with mitochondria absorbing calcium from the ER in response to EGF stimulation. A recent report showed that mitochondria localized to filopodia, termed METEORs, have a unique composition, enriched with MICOS, high levels of the Ca^2+^ uniporter MCU, and Myo19, suggesting a functional specialization. METEORs are also enriched for the myosin MYO19^63^. These findings indicate that mitochondria form a specialized subtype under cellular stress conditions, which is beneficial for calcium buffering and promotes mitochondrial translocation.

### Microtubule and Actin molecular motors coordinate mitochondrial trafficking

The localization of mitochondrial mass to the cell periphery and the tips of filopodia is a highly active and dynamic process. In this work, we demonstrate that this process is significantly enhanced by an integrated signaling pathway initiated via EGF stimulation, through calcium buffering that triggers mitochondrial motility, primarily involving two distinct motors, Kif5B and Myo19. We propose a five-step model for mitochondrial translocation. First, mitochondrial calcium uptake is followed by a fission event. Next, rapid anterograde transport of Kif5B along microtubules occurs, leading to its movement to the cell periphery. Then, spikes and waves of cytosolic calcium increase may regulate Kif5B’s activity via MIROS and TRACKS, by weakening its mitochondrial interactions or inhibiting Kif5B. At the same time, it remains bound, enabling Myo19 to become the dominant transporter and reach the actin filament projections of the cell^79,80^. Finally, Myo19 mediates the translocation of filopodia shafts toward their tips and helps coordinate mitochondrial backflow with actin filament dynamics and collapse within filopodia (Figure 7, 5-step model).

**Figure 7.**
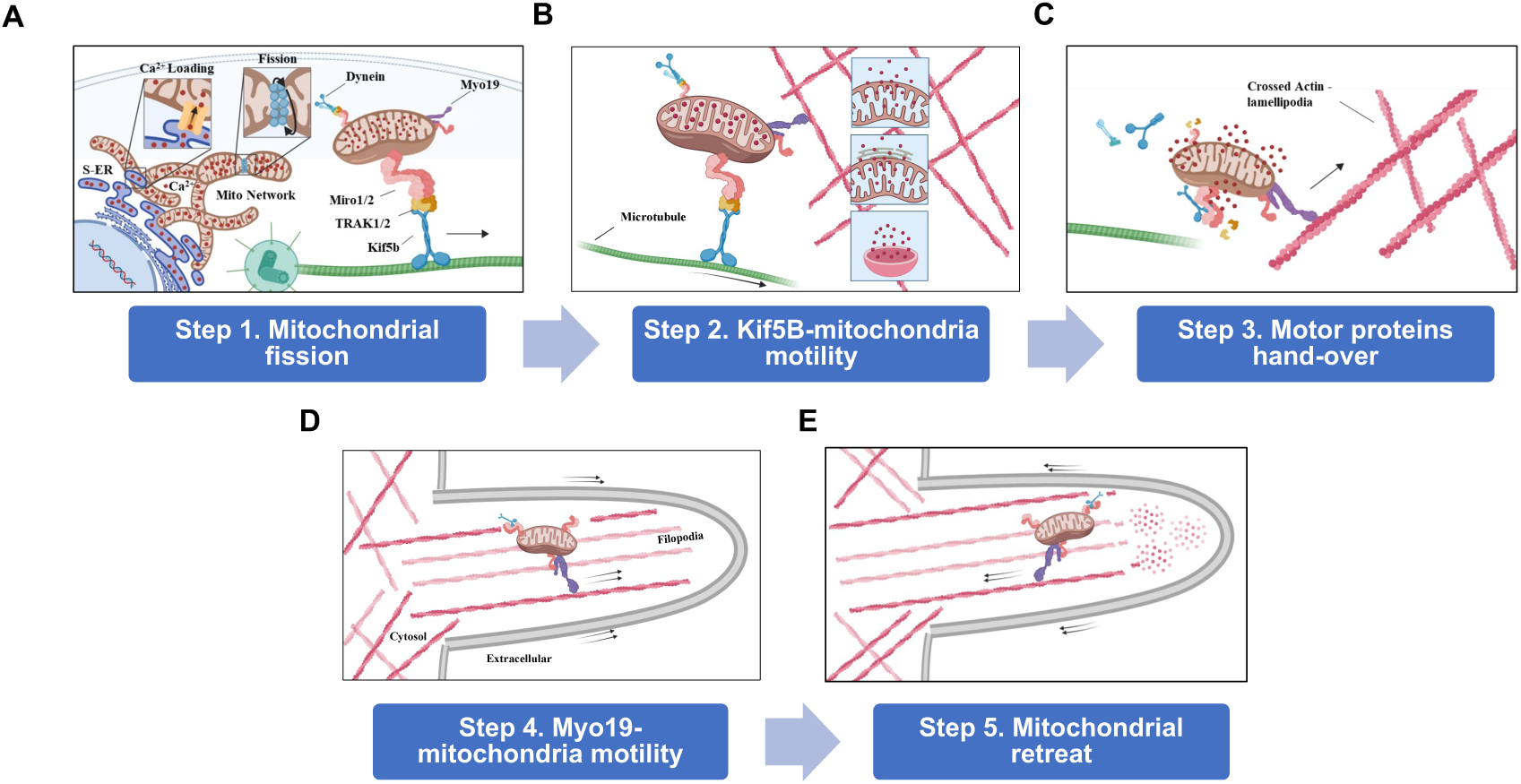
Recapitulation of mitochondrial translocation upon EGF stimulation. **(A)** Step 1: Mitochondrial calcium uptake from the ER and mitochondrial division. **(B)** Step 2: Kif5B translocases mitochondria to the cell’s leading edge, and mitochondria regulate calcium homeostasis. **(C)** Step 3: During the handover process, the concentration of cytosolic calcium increases, Kif5B dissociates from the mitochondria, and Myo19 regulates mitochondrial motility along actin filaments. **(D)** Step 4: Myo19-mitochondria translocated to the tips of filopodia. **(E)** Step 5: Mitochondrial retreat from filopodia is coupled with the dynamics of actin filaments.

### Unique features of Myo19-mitochondria-derived motility

Two notable features are observed in the velocity and directionality during the transfer of mitochondrial mass from the dense cell network to the periphery. There is a rapid phase that is more than twice as fast as in filopodia, where, in the actin-rich protrusion, the velocity is much slower and closely resembles the values seen in Myo19 in vitro motility actin gliding measurements^81,82^. This behavior, with similar velocities in cells and in vitro, was also reported for the motility of purified vesicles reconstituted in vitro, closely resembling the movement of LysoTracker-positive vesicles in primary neurons^83^. The processive bidirectional motility is interrupted by frequent directional switches, diffusional movement, and pauses due to the action of kinesin and dynein on the same vesicle. In our work on cells, we observe an overall unidirectional movement toward the cell periphery, characterized by dynamic motility within filopodia, which includes both extension and retraction. Overall, the motor system has specifically adapted to maintain the common properties shared by any vesicle cargo motility.

The second feature involves the alignment and splitting of mitochondria, making them thinner and aligned with the polarity of actin filaments at the base of the filopodia, followed by a fission event. We suggest that this process guides and facilitates a smooth transition from a dense, mixed cytoskeleton to actin-rich protrusions. Recent studies have shown that mitochondrial diameter ranges are under 100 nm in narrow cellular spaces, such as the TNT and the neuron axon^84,85^. Mitochondrial network remodeling in response to EGF stimulation indicates that variations in mitochondrial morphology can adapt to the specific functional space of the cell accordingly.

### Mitochondria motility bypasses barriers utilizing cytoskeletal motors

Sub-cellular organelle trafficking is a highly energy-consuming process that requires strict regulation and precise coordination. Organelles move along both types of cytoskeletal filaments using kinesins and myosins, which demands detailed cooperation, such as between mitochondrial motors^86^. Additionally, ER-mitochondria contact sites act as physical barriers that must be cleared to allow smaller mitochondrial masses to detach and be transported. However, positioning or detaching mitochondria across disconnected regions might help bypass this obstacle, making it easier to release their masses for transport toward the cell periphery. Our work focuses on organellar transport in non-neuronal cells, where the cytoskeletal network overlaps more extensively than in neuronal cell lines. More importantly, continuous tracks should allow switching between microtubules and actin filaments, thereby bridging the gaps between them. Therefore, we believe that network connectivity between microtubules and actin filaments is crucial for enabling full motility at the tips of cell protrusions. Ultimately, the cytoskeleton’s landscape likely influences the overall accuracy of navigation toward the base of newly formed filopodia.

### The kinesin-to-myosin switch may be simpler than in the case of tug-of-war

Biomechanical regulation of mitochondrial motors’ force generation depends on local concentrations of microtubule and actin, as well as the relative levels of ATP and ADP, along with the enzymatic adaptation of these molecular motors. The presence of alternative strong binding sites may provide a mechanism for track switching. Notably, the ATPase cycles of kinesin and myosin are fundamentally different, especially in their strong and dissociative states relative to their tracks. While unconventional myosins generally remain in a strongly bound state when ADP is attached, kinesins exhibit a weak binding state to microtubules in the presence of ADP. Conversely, at high ATP levels, kinesins bind strongly to microtubules, whereas myosins tend to dissociate^87–89^. Therefore, when mitochondria stall and a local drop in [ATP] occurs, Myo19 acts as a crosslinker, stopping instantly to increase [ATP] locally, as previously suggested^50^. As the Myo19-mitochondria complexes approach the cell periphery, actin filaments become denser and more projections, increasing the likelihood of Myo19 binding to an actin filament and initiating movement as a barbed-end Myo19 toward the base of lamellipodia and beyond to cellular protrusions after proper alignment and size fitting of the mitochondrial mass. Decreasing Kif5B motor density bound to mitochondria, as well as unloading calcium, may help adjust the mitochondria to fit within the limits of the filopodia actin tracks in the shaft.

### Molecular Motor Transport and Cellular Protrusion Dynamics

Theoretical models based on molecular motors, transport, and cellular protrusion dynamics have been rigorously reported for several myosin classes^60,90,91^. Pursuing a similar modeling approach for Myo19-mitochondria will be an essential component in linking its biomechanics to cellular physiology. However, because identical actin structures are utilized, it is most likely that the Myo19-mitochondria will use similar principles as previously suggested. These may include the distribution of Myo19-mitochondria across the filopodia and the motor dynamics partitioning between moderately processive movements, stalling, and diffusion during the collapse of protrusions. Finally, Myo19-mitochondria motility within the actin protrusions may be tightly coupled to their physical behavior, providing forces to regulate their dynamics and extensions.

This work provides a comprehensive understanding of how cells coordinate mitochondrial positioning during filopodia formation under physiological stress, particularly in cell lines highly relevant to their function during epidermal layer^92,93^formation. Ultimately, it reveals a fundamental mechanism of cellular energy distribution in processes such as migration and identifies potential therapeutic targets for cancer treatment.

### Limitations of the Study

Single-particle tracking of Kif5B was not possible due to its cellular distribution, which does not allow for monitoring Kif5B-mitochondria interactions at the temporal and spatial resolution necessary to measure its mobility directly.

## Methods

### Reagents

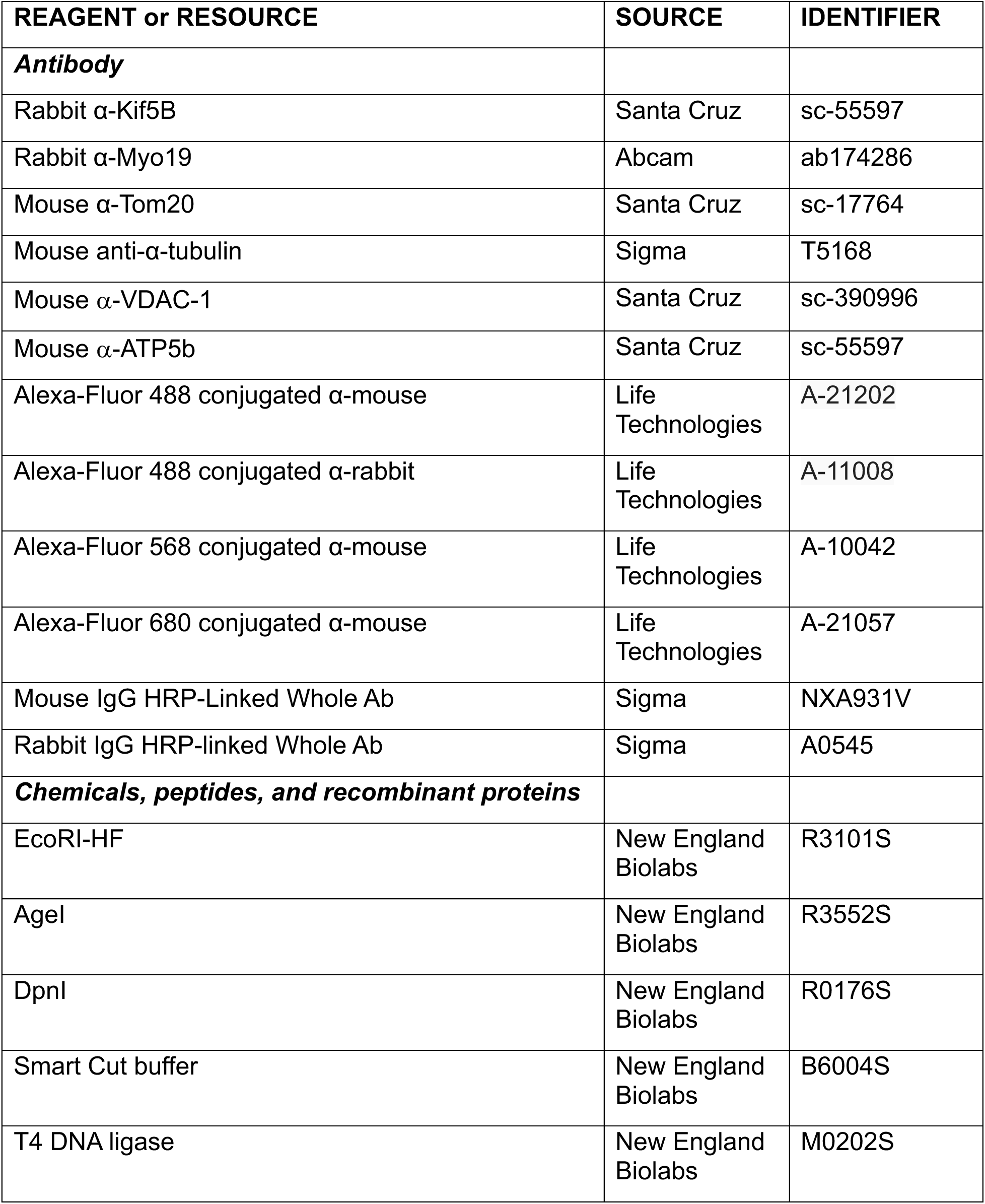
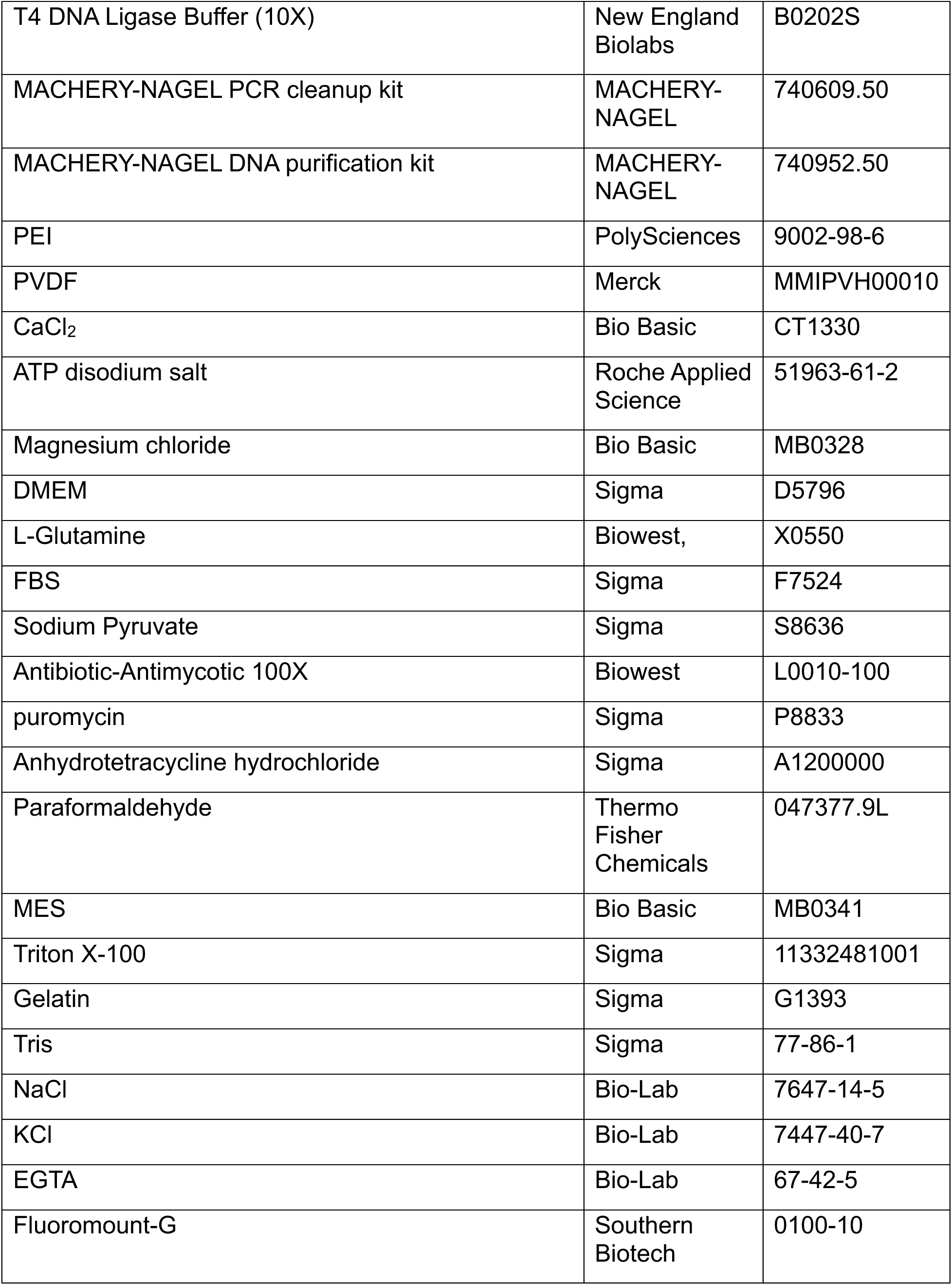
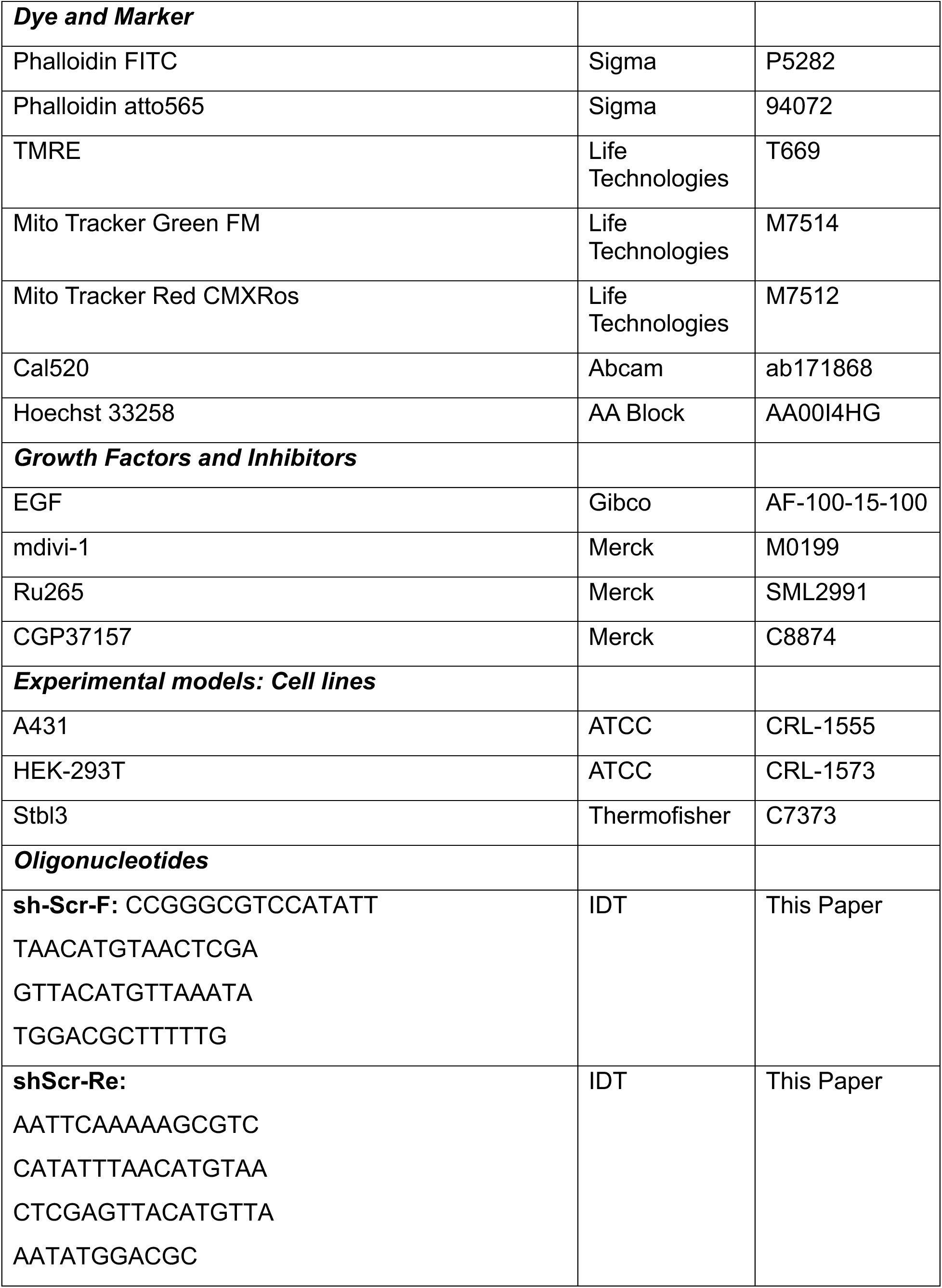
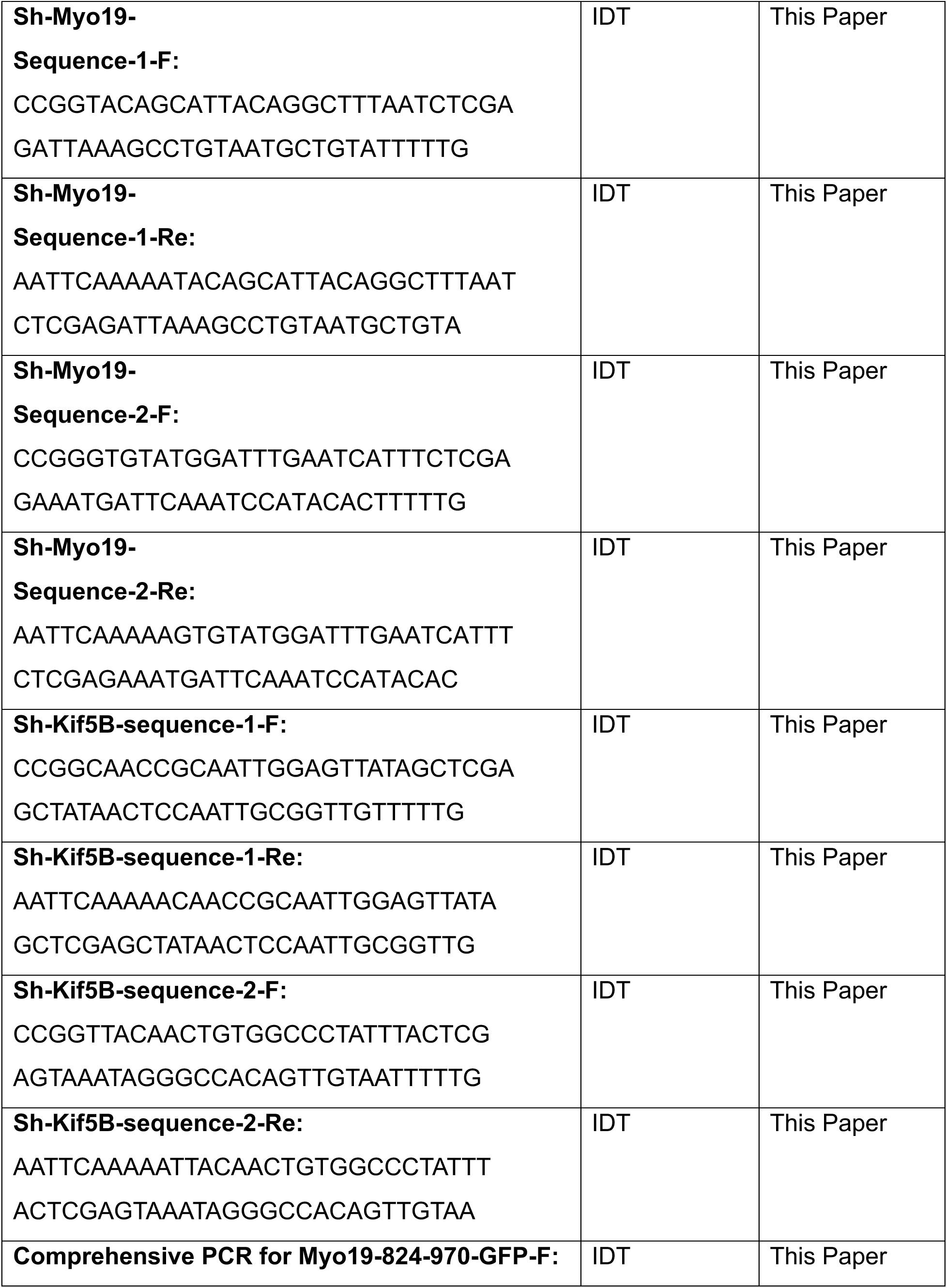
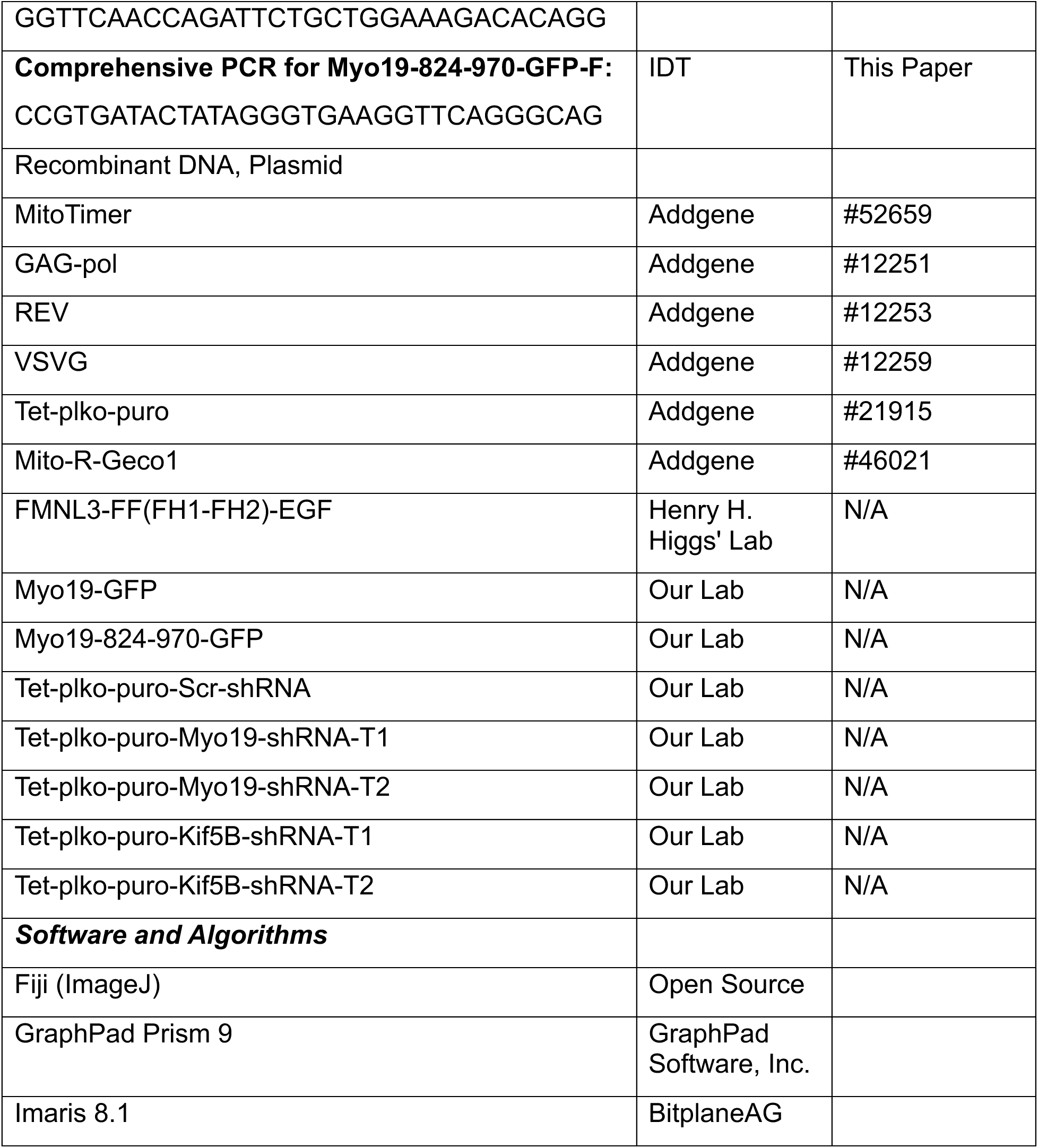

### Cell Lines

A431 human epidermoid carcinoma cells (ATCC) were cultured at 37°C with 5% CO₂ in Dulbecco’s modified Eagle’s medium (DMEM) supplemented with 10% fetal bovine serum (FBS), 4.5 g/L glucose, 4 mM L-glutamine, and 1 mM sodium pyruvate. Cell line authentication was performed by STR profiling, and mycoplasma testing was conducted regularly.

#### Molecular Cloning and Plasmid Construction

The Myo19-824-970-GFP construct containing a tail domain mutation was generated to enable expression in A431 cells following Myo19 knockdown. PCR amplification was performed using forward primer (5’-GGTTCAACCAGATTCTGCTGGAAAGACACAGG-3’) and reverse primer (5’-CCGTGATACTATAGGGTGAAGGTTCAGGGCAG-3’). The tail domain mutation did not alter the amino acid sequence of the Myo19 tail domain. Tet-inducible shRNA plasmids (Tet-plko-puro-Scr-shRNA, Tet-plko-puro-Myo19-shRNA-T1, Tet-plko-puro-Myo19-shRNA-T2, Tet-plko-puro-Kif5B-shRNA-T1, and Tet-plko-puro-Kif5B-shRNA-T2) were constructed using synthesized shRNA oligonucleotides (IDT) (sequences in Table S1). To construct the Tet-plko-puro-Scr-shRNA, the forward primer (5’-CCGGGCGTCCATATTTAACATGTAACTCGAGTTACATGTTAAATATGGACGCTTTT-TG-3’) and reverse primer (5’-AATTCAAAAAGCGTCCATATTTAACATGTAACTCGA-GTTACATGTTAAATATGGACGC-3’) were used to create annealed oligonucleotides. To construct the Tet-plko-puro-Myo19-shRNA-T1, the forward primer (5’-CCGGTACAGC- ATTACAGGCTTTAATCTCGAGATTAAAGCCTGTAATGCTGTATTTTTG-3’) and reverse primer (5’-CCGGTACAGCATTACAGGCTTTAATCTCGAGATTAAAGCCT-GTAATGCTGTATTTTTG-3’) were used to create annealed oligonucleotides. To construct the Tet-plko-puro-Myo19-shRNA-T2, the forward primer (5’-CCGGGTGTAT- GGATTTGAATCATTTCTCGAGAAATGATTCAAATCCATACACTTTTTG-3’) and reverse primer (5’-AATTCAAAAAGTGTATGGATTTGAATCATTTCTCGAGAAATGA-TTCAAATCCATACAC-3’) were used to create annealed oligonucleotides. To construct the Tet-plko-puro-Kif5B-shRNA-T1, the forward primer (5’-CCGGCAACCGCAATTGGA- GTTATAGCTCGAGCTATAACTCCAATTGCGGTTGTTTTTG-3’) and reverse primer (5’-AATTCAAAAACAACCGCAATTGGAGTTATAGCTCGAGCTATAACTCCAATTGCGGTTG-3’) were used to create annealed oligonucleotides. To construct the Tet-plko-puro-Kif5B-shRNA-T2, the forward primer (5’-CCGGTTACAACTGTGGCCCTATTTACTCG- AGTAAATAGGGCCACAGTTGTAATTTTTG-3’) and reverse primer (5’- AATTCAAAAATT- ACAACTGTGGCCCTATTTACTCGAGTAAATAGGGCCACAGTTGTAA -3’) were used to create annealed oligonucleotides.

The Tet-plko-puro vector (Addgene #21915) was linearized using EcoRI-HF and AgeI restriction enzymes in Smart Cut buffer overnight at 37°C. Following heat inactivation (20 min at 65°C), products were gel-purified using a MACHEREY-NAGEL PCR cleanup kit. Annealed oligonucleotides were ligated into the linearized vector using T4 ligase overnight at 16°C. Ligated plasmids were transformed into Stbl3 (Thermofisher, C7373)- competent cells, and positive clones were verified by Sanger sequencing.

#### Lentiviral Production and Transduction

Lentiviral particles were produced by co-transfecting HEK-293T cells (ATCC, CRL-1573) with four packaging plasmids: GAG-pol (Addgene, #12251), REV (Addgene, #12253), VSVG (Addgene, #12259), and the respective Tet-Plko-Puro shRNA construct using polyethylenimine (PEI, Polysciences). Plasmid DNA and PEI were separately diluted in 150 mM NaCl, combined, and incubated for 25 minutes at room temperature before being added to the cells. Viral supernatants were collected at 48 and 72 h post-transfection, filtered through 0.45 μm cellulose acetate filters, and stored at −80°C.

For transduction, A431 cells were seeded at 0.5×10⁶ cells per well in 6-well plates. Lentiviral supernatant (1 mL) was added twice daily at 12 h intervals for three consecutive days in the presence of 1 μg/mL polybrene. Transduced cells were selected with 0.8 μg/mL puromycin (concentration determined by kill curve analysis over 10 days).

#### Single Cell Cloning and shRNA Induction

A431 cells were trypsinized, resuspended at ≤ 2×10⁶ cells/mL, and filtered through 35 μm strainers. Single cells were sorted into 96-well plates using a BIGFOOT sorter (Invitrogen) based on forward and side scatter parameters at a flow rate of 0.3 PSI. Cells were cultured in DMEM supplemented with 4.5 g/L glucose, 20% FBS, 1× antibiotics, and 1 mM sodium pyruvate. Five monoclonal colonies were screened for knockdown efficiency by western blotting.

For shRNA induction, cells were seeded at 5×10⁴ cells/well in 6-well plates and treated with 1 μg/mL anhydrotetracycline hydrochloride (Atet-HCl; Sigma A1200000). The medium containing fresh Atet-HCl was replaced every 24 hours for 96 hours. Whole cell lysates were prepared for Western blot analysis.

#### Serum Starvation and EGF stimulation

Cells were washed three times with PBS and incubated in starvation medium (DMEM with 4.5 g/L glucose, 0.1% FBS, antibiotics, and 1 mM sodium pyruvate) for 16 h. For EGF stimulation, cells were treated with 30 ng/mL EGF for 30 minutes.

For inhibitor treatments: (1) Drp1 inhibition - cells were pretreated with 50 μM mdivi-1 for 6 h, then stimulated with EGF in the presence of mdivi-1 for 30 min. (2) MCU and NCLX inhibition - cells were pretreated with 1 μM Ru265 or 10 μM CGP37157 for 10 min, then stimulated with EGF in the presence of inhibitors for 30 min.

#### Live Cell Imaging

For live imaging experiments, cells were serum-starved for 15 h. They were transferred to 35 mm optical plates containing 1.8 mL of Live Imaging Medium (DMEM with 0.1% FBS, antibiotics, and 5 mM sodium pyruvate). The plates were equilibrated in a microscope incubator (37°C, 5% CO₂) for 1 h before adding 200 μL of DMEM with 30 ng/mL EGF (final concentration). For inhibitor experiments, compounds were applied 10 minutes before EGF stimulation.

To track Myo19-mitochondrial motility, express Myo19-GFP in A431 cells using the Myo19-GFP plasmid (Our lab), and stain the mitochondria with TMRE (Life Technologies, T669). Mitochondrial membrane potential was measured by the ratio of Mito Tracker Green FM (Life Technologies, M7514) to TMRE fluorescence intensity. A431 cells were electroporated with MitoTimer (Addgene, #52659), a marker for mitochondrial health, as visualized by live imaging. The FMNL3-FF(FH1-FH2)-GFP plasmid (Henry H. Higgs’ Lab) was electroporated into A431 cells to track FMNL3-specific filopodia elongation upon EGF stimulation. Cal520 (Abcam, Ab171868) was used to measure calcium concentration in the cytoplasm. Mito-R-Geco1(Addgene, #46021) was electroporated into A431 cells, which is a marker for mitochondrial calcium measurement. Hoechst 33258 (AA Block, AA0014HG) was used to stain the nucleus.

#### Immunofluorescence Microscopy

Cells grown on coverslips were fixed with 4% paraformaldehyde (Thermo Fisher Chemicals, 047377.9L) in CBS buffer (10 mM MES pH 6.1, 138 mM KCl, 3 mM MgCl₂, 2 mM EGTA, 7.8% sucrose) for 10 min. After permeabilization and blocking with 10% BSA in TBST (0.3% Triton X-100 in Tris-buffered saline) for 1 h, cells were incubated with primary antibodies: Rabbit α-Kif5B (Santa Cruz, sc-55597), Rabbit α-Myo19 (Abcam, ab174286), Mouse α-Tom20 (Santa Cruz, sc-17764), Mouse anti-α-tubulin(Sigma, T5168), overnight at 4°C, followed by fluorescent secondary antibodies: Alexa-Fluor 488 conjugated α-mouse (Life Technologies, A-21202), Alexa-Fluor 488 conjugated α-rabbit (Life Technologies, A-11008), Alexa-Fluor 568 conjugated α-mouse (Life Technologies, A10042), Alexa-Fluor 680 conjugated α-mouse (Life Technologies, A21057), for 1 h at room temperature. F-actin was visualized using fluorescent phalloidin: Phalloidin FITC (Sigma, P5282) or Phalloidin atto565 (Sigma, 94072). Nuclei were counterstained with Hoechst 33342 (AA Block, AA0014HG), and coverslips were mounted using Fluoromount-G (Southern Biotech, 0100-10). The Mito Tracker Red CMXRos (Life Technologies, M7512) was used to stain mitochondria before fixation.

#### Microscopy and Image Acquisition

Fixed samples were imaged using an LSM700 confocal microscope (Zeiss) with a 63× oil immersion objective (pixel size ∼0.076 μm, 8-bit depth) at 37°C. Excitation wavelengths were 488, 543, and 639 nm.

For motor protein-mitochondria colocalization analysis, images were captured using an LSM980 microscope (Zeiss) with Airyscan detection. Kif5B-mitochondria samples were imaged as 5 z-stack slices (150 nm apart) and processed with Airyscan2 3D. Myo19-mitochondria samples were imaged as 3 z-stack slices (150 nm apart) and processed with Airyscan2 2D.

Live imaging was performed using an LSM710 microscope (mitochondrial motility) or Nikon Spinning Disk Confocal (Ca²⁺ measurements) at 37°C. For mitochondrial dynamics, images were acquired at 1-second intervals following EGF stimulation. For Ca²⁺ measurements, images were captured starting 10 seconds after EGF addition at 10-second intervals.

#### Electroporation

Cells were electroporated using the Nucleofector X Unit (Lonza). Solution A (0.2 g/mL ATP disodium salt and 0.12 g/mL MgCl₂; 80 μL) was combined with 4 mL of Solution B. Cell pellets were resuspended in 100 μL of the combined solution, mixed with DNA, and transferred to electroporation cuvettes. After electroporation using the A431 cell program, cells were immediately transferred to pre-warmed growth medium.

#### Western Blot Analysis

Cell lysates were prepared and subjected to SDS-PAGE. Proteins were transferred to PVDF membranes using the eBlot L1 Fast Wet Transfer System (GeneScript) for 26 min. Membranes were blocked with 5% milk powder for 1 h at room temperature, incubated with primary antibodies: Rabbit α-Kif5B (Santa Cruz, sc-55597), Rabbit α-Myo19 (Abcam, ab174286), Mouse α-VDAC-1 (Santa Cruz, sc-390996), Mouse α-ATP5b (Santa Cruz, sc-55597), overnight at 4°C, and detected using HRP-conjugated secondary antibodies: Mouse IgG HRP Linked Whole Ab (Sigma, NXA931V), Rabbit IgG HRP linked Whole Ab (Sigma, A0545). Chemiluminescent signals were developed using a luminol-based substrate and imaged using a VILBER Fusion 400 system.

#### Image Analysis and Quantification

Images were processed using ImageJ software (Open Source). Filopodia length and number were quantified using the FiloQuant plugin^94^. Mitochondrial dynamics were manually classified, and motor protein-mitochondria colocalization was analyzed using the JACop plugin^95^ to calculate Pearson correlation coefficients. Mitochondrial velocity tracking was performed using Imaris 8.1 software (BitplaneAG), which utilizes surface analysis and autoregressive motion tracking algorithms.

#### Quantification and Statistical Analysis

Statistical analyses were performed using GraphPad Prism version 9 (GraphPad Software, Inc.). Data are presented as mean ± SEM from at least three independent experiments. The statistical significance between the two groups was determined using an unpaired Student’s t-test. For multiple group comparisons, one-way ANOVA followed by Tukey’s multiple comparisons test was used. Contingency data for mitochondrial dynamics were analyzed using chi-square tests. Mitochondrial velocity distributions were analyzed using nonlinear regression with Gaussian distribution models. Statistical significance was defined as *p < 0.05, **p < 0.01, ***p < 0.001. Sample sizes (n) are indicated in figure legends.

## Supporting information

SLMyo19CaSuppFigures

SLMyo19CaSuppTables

## RESOURCE AVAILABILITY

### Lead contact

Further information and requests for resources and reagents should be directed to and will be fulfilled by the lead contact, Arnon Hen

### Materials Availability

Plasmids generated in this study are available from the lead contact upon request.

### Data and Code Availability

Data reported in this paper will be shared by the lead contact upon request. This paper does not report original code.

## Acknowledgments

This study was supported by the Israel Scientific Foundation (ISF grant # 712/18). We thank Drs. Atan Gross and Nir S. Gov for their valuable comments and critical reading of the manuscripts. We thank Dr. Nitsan Dahan for his support in the microscopy work presented in this work.

## Author Contributions

Conceptualization by SL., B.I.S., and A.H. Methodology by SL., B.I.S., and A.H. Data analysis by S.L. and A.H. Drafting and editing by S.L. and A.H. All authors revised and reviewed the paper and approved the final version.

## Supplemental Information

Supplement Figures S1-S6

Supplement Movies S1-S6

Supplement Table S1

Supplement Table S2

