## Supplementary material for "Calcium-Regulated Mitochondria Remodeling by Myo19 is Required for Filopodia Tip-Extension": SLMyo19CaSuppFigures

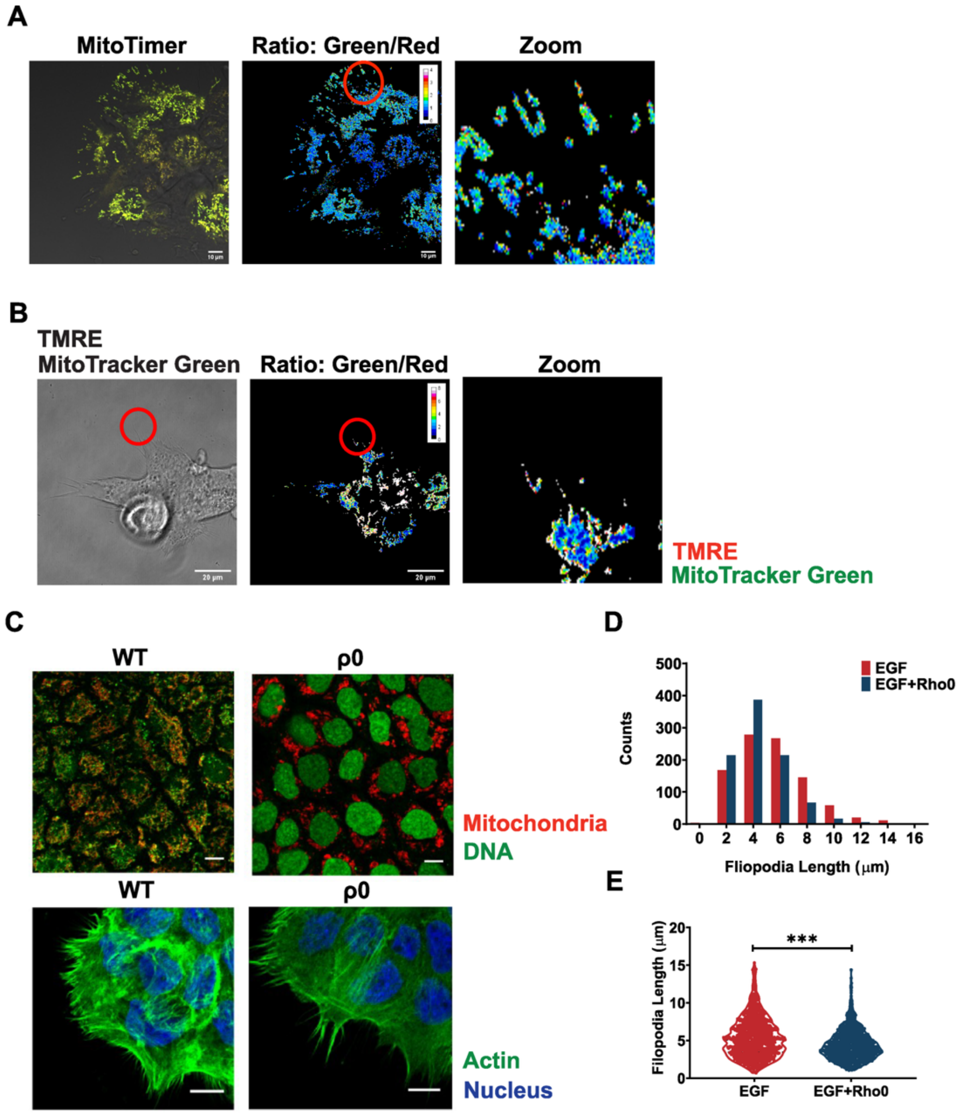

A
MitoTimer
Ratio: Green/Red
Zoom
B
TMRE
MitoTracker Green
Ratio: Green/Red
Zoom
TMRE
MitoTracker Green
C
WT
ρ0
Mitochondria
DNA
WT
ρ0
Actin
Nucleus
D
EGF
EGF+Rho0
Counts
Fliopodia Length (μm)
E
***
Filopodia Length (μm)
EGF
EGF+Rho0

**Figure S1. Mitochondrial aging, membrane potential, and mtDNA-dependent mitochondrial behavior in EGF stimulated filopodia.**

**(A)** Ratiometric imaging of EGF-stimulated A431 stable cell lines expressing MitoTimer. Upon oxidation, MitoTimer fluorescence shifts from green (excitation/emission: 488/518 nm) to red (excitation/emission: 543/572 nm), as shown in the ratio image and zoomed view, revealing mitochondrial aging patterns. The scale bars are 10 μm.

**(B)** Ratiometric imaging of EGF-stimulated wildtype (WT) A431 cells co-stained with TMRE (membrane potential-sensitive, red) and MitoTracker Green (membrane potential-insensitive, green). Red circles indicate regions of interest. The scale bars are 25 μm.

**(C)** Representative images of WT and ρ0 A431 cells stained with PicoGreen demonstrating mtDNA depletion in ρ0 cells (upper panels). Lower panels show phalloidin-stained actin filaments (green), mitochondria (red), and DAPI-stained nuclei (blue) in WT and ρ0 cells. The scale bars are 10 μm.

**(D)** Histogram showing distribution of filopodia length in EGF-stimulated WT and ρ0 A431 conditions.

**(E)** Violin plots comparing filopodia length in EGF-stimulated WT(red) and ρ0 A431 (blue) conditions. Data represent mean ± SEM (n ≥ 900 filopodia). ***$p < 0.001$ by unpaired t-test.

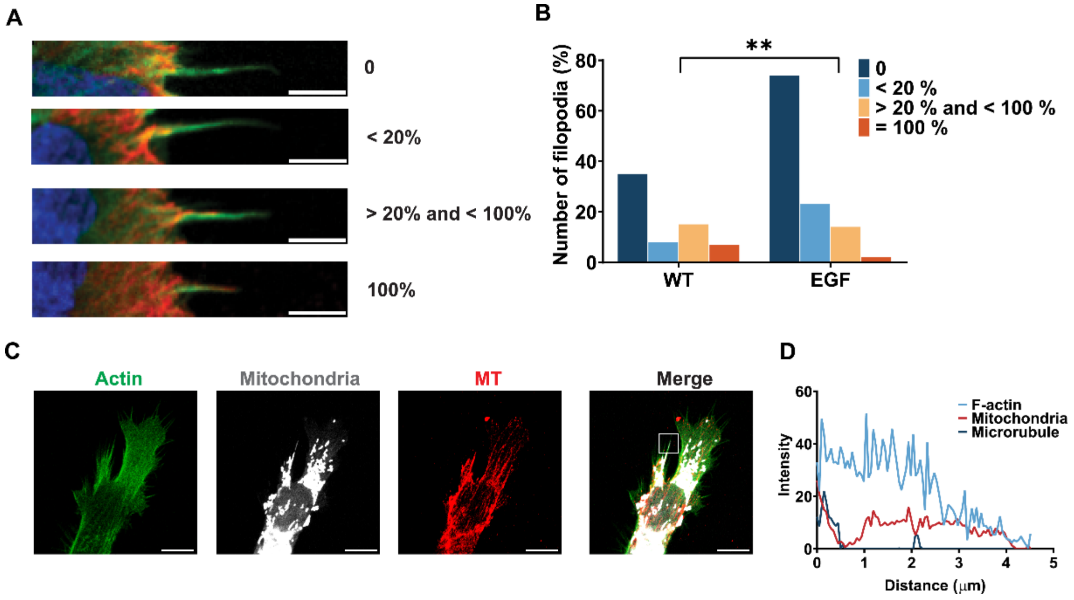


**Figure S2. Mitochondrial distribution correlates with cytoskeletal architecture in filopodia.**

**(A)** Representative confocal fluorescence microscopy images showing microtubule and actin filament distribution within filopodia. Categories indicate microtubule-to-actin length ratio: 0 (microtubules absent), <20% (microtubules occupy <20% of filopodia length), >20% and <100% (microtubules occupy 20-100% of filopodia length), and 100% (microtubules span entire filopodia). Scale bar, 10 μm.

**(B)** Quantitative analysis of MT and actin filament length ratios in filopodia from untreated (WT) and EGF-stimulated cells. EGF treatment significantly increases the proportion of filopodia with absent or minimal microtubule content (0 category) while decreasing the proportion of filopodia with extensive microtubule networks ($n \geq 60$ filopodia). $^{**}p < 0.01$ by chi-square test.

**(C)** Confocal fluorescence microscopy showing mitochondrial localization in filopodia containing predominantly actin filaments with minimal MT content. F-actin (green), mitochondria (white), microtubules (red), and merged images are displayed. Scale bar, 10 μm.

**(D)** Fluorescence intensity profile along filopodia length corresponding to the boxed region in (C), showing distribution of F-actin (blue), mitochondria (red), and microtubules (black). The profile shows mitochondrial positioning within actin-rich regions with minimal overlap of microtubules.

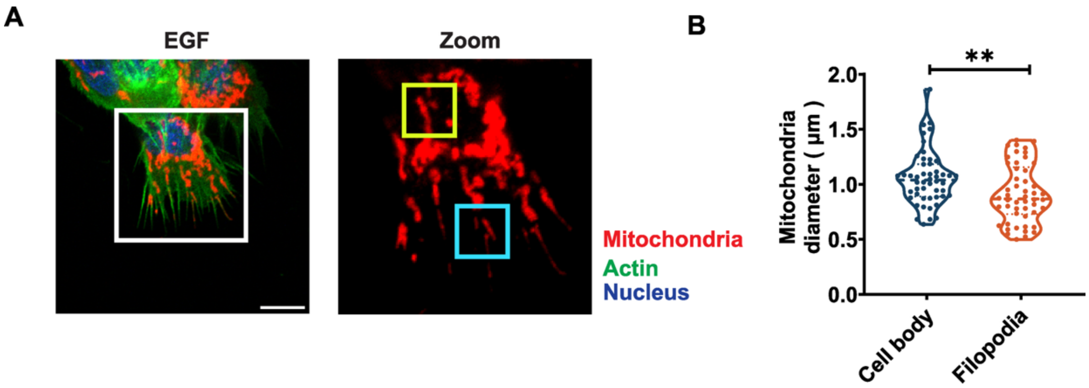


**Figure S3. The diameter of mitochondria in the cell body and filopodia upon EGF stimulation.**

**(A)** Confocal fluorescence microscopy of EGF-stimulated A431 cells showing F-actin (green), mitochondria (red), and nuclei (blue). The left panel displays an overview with a white box indicating the region that will be zoomed in on. The right panel shows a magnified view with a yellow box highlighting mitochondria in the cell body and a cyan box indicating mitochondria within filopodia. Scale bar is 10 μm.

**(B)** Quantification of mitochondrial diameter in the cell body versus filopodia. Mitochondria in filopodia exhibit significantly smaller diameters compared to those in the cell body (n = 60 mitochondria per location). Data presented as violin plots with individual data points. Statistical significance determined by unpaired t-test; **$p < 0.01$.

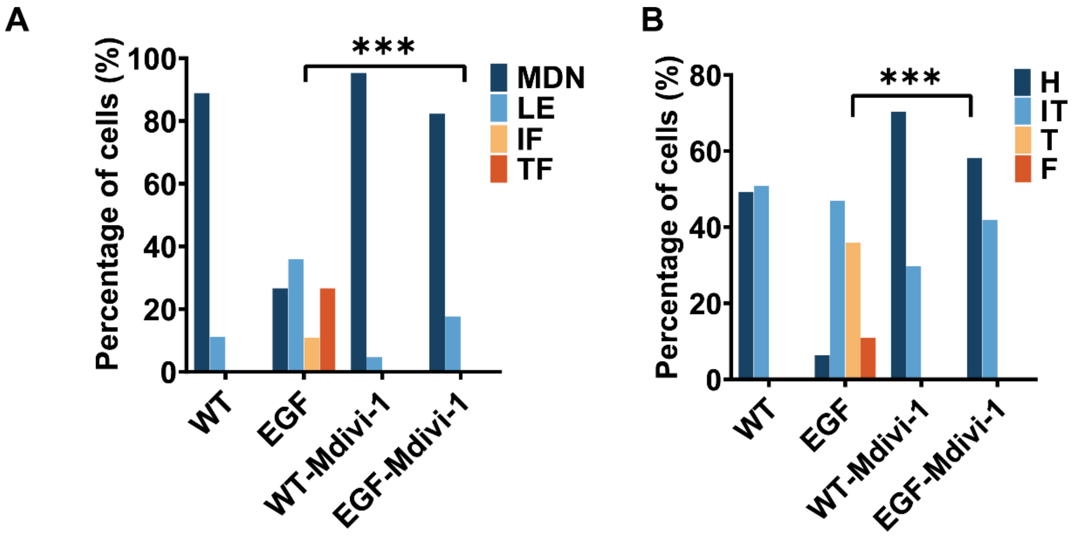


**Figure S4. Mitochondrial morphology and cellular localization during Drp1 inhibition.**

**(A and B)** Quantitative analysis of mitochondrial morphology **(A)** and subcellular localization **(B)** in A431 cells following Drp1 inhibition. Classification categories for mitochondrial morphology and localization are consistent with those shown in Figure 1. Data from three independent experiments ($n \geq 60$ cells per condition).

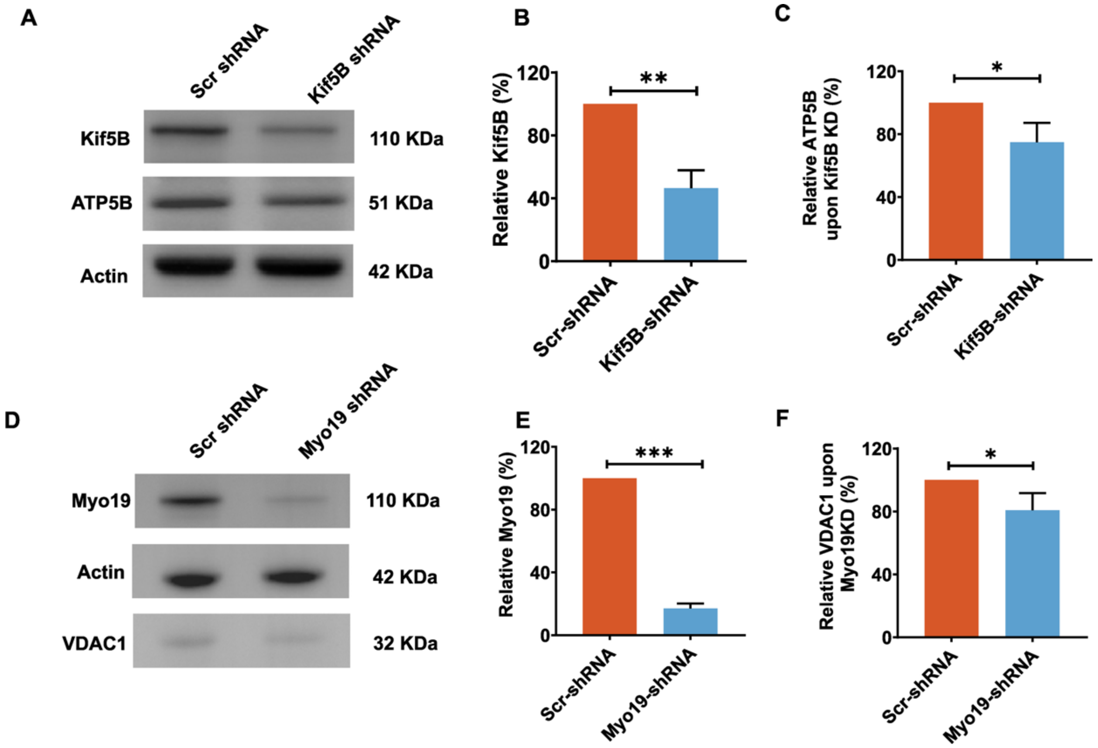


**Figure S5. Validation of Myo19 and Kif5B knockdown efficiency in A431 cells expressing inducible shRNA.**

**(A and B)** Western blot analysis of ATP5b expression in scrambled control (Scr) and Kif5B knockdown (KD) A431 cells. **(A)** Representative Western blot. **(B)** Quantification of ATP5b expression levels (n = 3). Data represent mean ± SEM. *$p < 0.05$ by unpaired t-test.

**(C and D)** Western blot analysis of VDAC1 expression in scrambled control and Myo19 knockdown (KD) A431 cells. **(C)** Representative Western blot. **(D)** Quantification of VDAC1 expression levels (n = 3). Data represent mean ± SEM. *$p < 0.05$ by unpaired t-test.

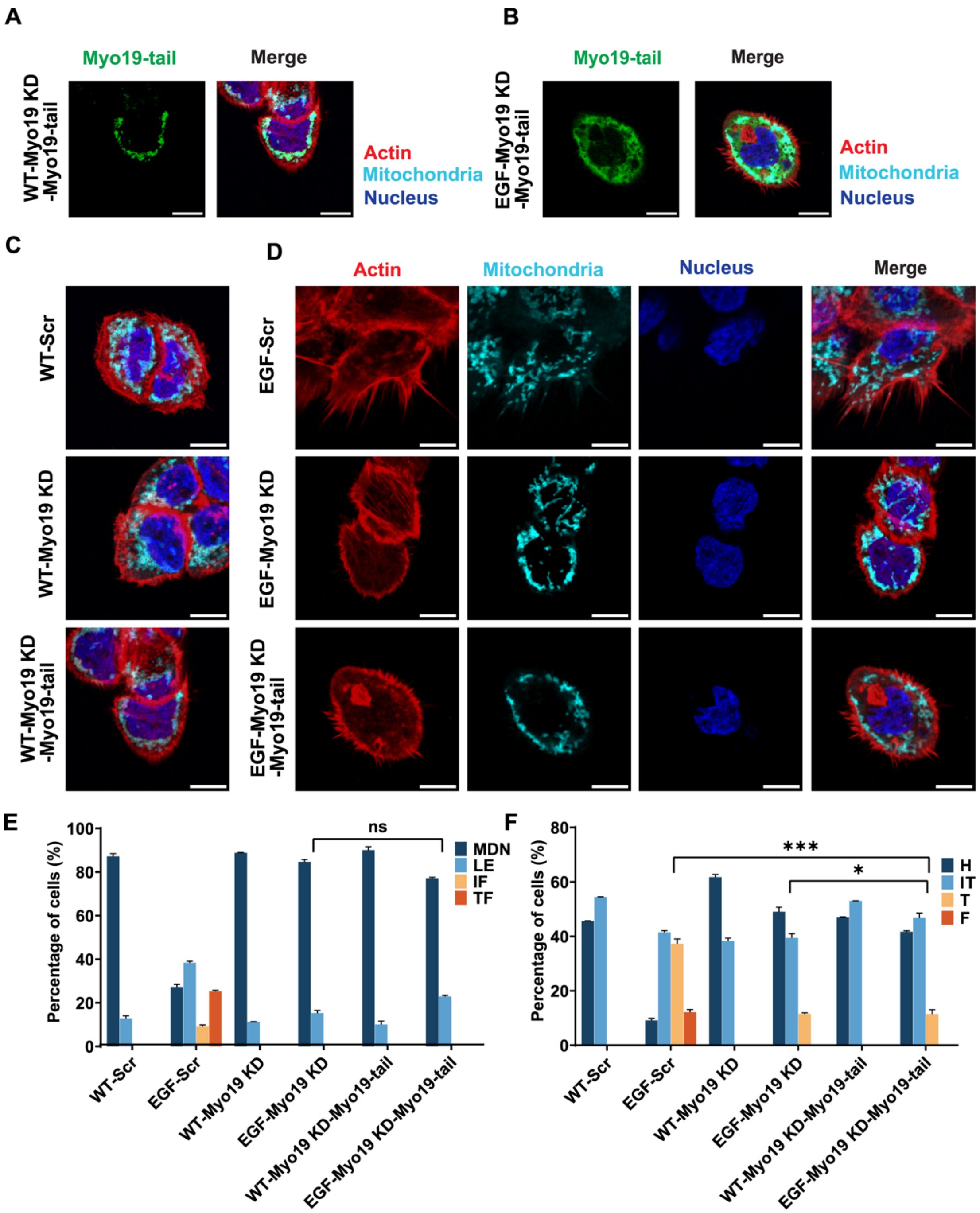
A
Myo19-tail
Merge
WT-Myo19 KD -Myo19-tail
Actin
Mitochondria
Nucleus
B
Myo19-tail
Merge
EGF-Myo19 KD -Myo19-tail
Actin
Mitochondria
Nucleus
C
WT-Scr
WT-Myo19 KD
WT-Myo19 KD -Myo19-tail
D
Actin
Mitochondria
Nucleus
Merge
EGF-Scr
EGF-Myo19 KD
EGF-Myo19 KD -Myo19-tail
E
Percentage of cells (%)
ns
MDN
LE
IF
TF
WT-Scr
EGF-Scr
WT-Myo19 KD
EGF-Myo19 KD
WT-Myo19 KD-Myo19-tail
EGF-Myo19 KD-Myo19-tail
F
Percentage of cells (%)
***
*
H
IT
T
F
WT-Scr
EGF-Scr
WT-Myo19 KD
EGF-Myo19 KD
WT-Myo19 KD-Myo19-tail
EGF-Myo19 KD-Myo19-tail

**Figure S6. Rescue experiments in Myo19 knockdown cells with Myo19-tail domain expression in A431 cells.**

**(A)** Confocal fluorescence microscopy of A431 cells expressing Myo19-tail following EGF stimulation. Myo19-tail (Green), actin filaments (red), and nuclei (blue). Scale bar, 10 μm.

**(B and C)** Confocal fluorescence microscopy showing mitochondrial localization and actin filaments in scrambled control (Scr), Myo19 knockdown (Myo19 KD), and Myo19 knockdown with Myo19-tail rescue (Myo19 KD-Myo19-tail) A431 cells. **(B)** Cells without EGF stimulation. **(C)** Cells with EGF stimulation. Scale bar, 10 μm.

**(D and E)** Quantitative analysis of mitochondrial morphology **(D)** and subcellular localization **(E)** across experimental conditions. Groups represent: WT/EGF (scrambled shRNA-expressing A431 cells), WT-/EGF-Myo19 KD (Myo19 knockdown shRNA-expressing A431 cells), and WT-/EGF-Myo19 KD-Myo19-tail (A431 cells co-expressing Myo19-tail and Myo19 knockdown shRNA). Classification categories for mitochondrial morphology and localization are consistent with those shown in Figure 1. Data from three independent experiments ($n \geq 90$ cells per condition).
