## Supplementary material for "Calcium-Regulated Mitochondria Remodeling by Myo19 is Required for Filopodia Tip-Extension": SLMyo19CaSuppTables

Table *S*1: *Mitochondria translocate to filopodia upon EGF stimulation*

Table S1A: Figure 1B

| Filopodia length quantification | |
|---|---|
| Cell Group | Filopodia Length (µm) |
| wildtype | 0.9 ± 0.1 |
| EGF stimulation | 1.9 ± 0.1 |

Table S1B: Figure 1C

| Number of filopodia per cell | |
|---|---|
| Cell Group | Number of Filopodia (counts) |
| wildtype | 1.6 ± 0.2 |
| EGF stimulation | 11.9 ± 0.5 |

Table S1C: Figure 1E

| Mitochondrial positioning (%) | | | | |
|---|---|---|---|---|
| Cell Group | MDN | LE | IF | TF |
| wildtype | 87.7 | 12.3 | 0 | 0 |
| EGF stimulation | 27.5 | 37.9 | 7.1 | 27.5 |

Table S1D: Figure 1G

| Mitochondrial morphology (%) | | | | |
|---|---|---|---|---|
| Cell Group | Hyperfused | Intermediate Tubular | Tubular | Fragmented |
| wildtype | 46.7 | 53.3 | 0 | 0 |
| EGF stimulation | 10.3 | 40.8 | 37.0 | 11.9 |

Table S1E: Figure S1

| Mitochondrial diameter | |
|---|---|
| Cell Group | Diameter (µm) |
| Cell body | 1.0 ± 0.2 |
| Filopodia | 0.9 ± 0.2 |

Table S1F: Figure S2B

| MT occupancy (%) | | | | |
|---|---|---|---|---|
| Cell Group | 0 | < 20% | 20% > and <100% | 100% |
| wildtype | 53.8 | 12.3 | 23.1 | 10.8 |
| EGF stimulation | 65.5 | 20.3 | 12.4 | 1.8 |

TableS1G: Figure S3D

| Filopodia length | |
|---|---|
| Cell Group | Length (μm) |
| EGF stimulation | 5.5 ± 0.1 |
| EGFs+Rho0 | 4.5 ± 0.1 |

Table S2*: Mitochondria fission affects mitochondria translocation upon EGF stimulation*

Table S2A: Figure 2B

| Filopodia length quantification | |
|---|---|
| Cell Group | Filopodia Length (μm) |
| wildtype | 1.0 ± 0.1 |
| EGF stimulation | 2.3 ± 0.1 |
| WT-mdivi-1 | 0.8 ± 0.1 |
| EGF stimulation -mdivi-1 | 0.9 ± 0.1 |

Table S2B: Figure 2C

| Number of filopodia per cell | |
|---|---|
| Cell Group | Number of Filopodia (counts) |
| wildtype | 1.6 ± 0.2 |
| EGF stimulation | 10.8 ± 0.7 |
| WT-mdivi-1 | 0.9 ± 0.2 |
| EGF stimulation -mdivi-1 | 1.3 ± 0.2 |

Table S2C: Figure S4A

| Mitochondrial positioning (%) | | | | |
|---|---|---|---|---|
| Cell Group | MDN | LE | IF | TF |
| wildtype | 88.9 | 11.1 | 0 | 0 |
| EGF stimulation | 26.6 | 35.9 | 10.9 | 26.6 |
| WT-mdivi-1 | 95.3 | 4.7 | 0 | 0 |
| EGF stimulation -mdivi-1 | 82.3 | 17.7 | 0 | 0 |

Table S2D: Figure S4B

| Mitochondrial morphology (%) | | | | |
|---|---|---|---|---|
| Cell Group | Hyperfused | Intermediate Tubular | Tubular | Fragmented |
| wildtype | 49.2 | 50.8 | 0 | 0 |
| EGF stimulation | 6.3 | 46.9 | 35.9 | 10.9 |
| WT-mdivi-1 | 70.3 | 29.7 | 0 | 0 |
| EGF stimulation -mdivi-1 | 58.1 | 41.9 | 0 | 0 |
